# Heterogeneous, temporally consistent, and plastic brain development after preterm birth

**DOI:** 10.1101/2024.12.06.627134

**Authors:** Melissa Thalhammer, Jakob Seidlitz, Antonia Neubauer, Aurore Menegaux, Benita Schmitz-Koep, Maria A. Di Biase, Julia Schulz, Lena Dorfschmidt, Richard A. I. Bethlehem, Aaron Alexander-Bloch, Chris Adamson, Gareth Ball, Claus Zimmer, Marcel Daamen, Henning Boecker, Peter Bartmann, Dieter Wolke, Dennis M. Hedderich, Christian Sorg

## Abstract

The current view of neurodevelopment after preterm birth presents a strong paradox: diverse neurocognitive outcomes suggest heterogeneous neurodevelopment, yet numerous brain imaging studies focusing on average dysmaturation imply largely uniform aberrations across individuals^1^. Here we show both, spatially heterogeneous individual brain abnormality patterns (IBAPs) but with consistent underlying biological mechanisms of injury and plasticity. Using cross-sectional structural magnetic resonance imaging data from preterm neonates and longitudinal data from preterm children and adults in a normative reference framework^2^, we demonstrate that brain development after preterm birth is highly heterogeneous in both severity and patterns of deviations. Individual brain abnormalities were also consistent for their extent and location along the life course, associated with glial cell underpinnings, and plastic for influences of the early social environment. Thus, IBAPs of preterm birth are spatially heterogenous, temporally consistent for extent, spatial location, and cellular underpinnings, and plastic for social-environmental impacts. Our findings extend conventional views of preterm neurodevelopment, revealing a nuanced landscape of individual variation, with consistent commonalities between subjects. This integrated perspective of preterm neurodevelopment implies more targeted theranostic intervention strategies, specifically integrating brain charts^2^ and imaging at birth, as well as social interventions during early development^3^.

## Main

In humans, preterm birth is defined as birth before 37 weeks of gestation^4^. With a worldwide prevalence of about 11 %^5^, it is the leading cause of perinatal mortality and long-term motor, cognitive, and behavioral impairments^6–8^. Impairments are typically more severe with decreased gestational age (GA) and lower quality of postnatal care^9,10^. Preterm birth impacts brain development through initial brain injuries that result in dysmaturation patterns affecting multiple tissue types and brain regions^1,11^. Initial injuries like hypoxic-ischemic events damage vulnerable cell populations, mainly the pre-oligodendrocyte (OL) cell line and the subplate, with subsequent local inflammation mediated by reactive astrocytes and activated microglia^11–17^. Resulting dysmaturation patterns of preterm birth have been shown to affect most parts of the brain, i.e., widespread gray matter areas, including cortical regions as well as basal ganglia, amygdala, thalamus, and hypothalamus nuclei^18–26^, and white matter areas^27–32^, not only in infants and children, but also in adolescents and adults. For example, cortical thickness (CTh) reductions might reflect various microscopic processes affected by prematurity, such as accelerated white matter maturation^33,34^. Group average focused brain magnetic resonance imaging (MRI) studies suggest that CTh is persistently altered in preterm-born subjects compared to full-term controls, with widespread CTh increases in infancy, a faster thinning rate in adolescence, and widespread decreases in adulthood^32,35–38^. These findings contribute to a model of widespread altered brain development following preterm birth largely shared between individuals, explaining injury-induced patterns of average dysmaturation outcomes at different stages of development^1,39,40^.

Whereas this model adequately describes the average brain aberrations of prematurity, it disregards, however, potential individual heterogeneity of altered brain development. Heterogeneity following preterm birth is evident for several outcomes^8,41–45^, including postmortem neonatal brain aberrations as well as neurocognitive functioning. In particular, the average IQ after very preterm birth (i.e., before 32 weeks of gestation) is about 11 points lower than that of full-term peers in childhood and adulthood, but individual IQ varies considerably between subjects^41,46–48^. This neurocognitive heterogeneity is widely assumed to be reflected in related heterogeneity of neurodevelopmental processes^1,49,50^. Thus, considering vast heterogeneity in neurocognitive-behavioral outcomes alongside persistent and widespread average brain aberrations, we hypothesized substantial spatial heterogeneity of individual brain abnormality patterns (IBAPs) after preterm birth across development. Despite suspected heterogeneity, we expected consistency in specific aspects of IBAPs, particularly concerning features related to injury-induced dysmaturation and developmental plasticity. Regarding injury-induced dysmaturation, we hypothesized a consistent impact of initial injury on IBAPs in three domains: extent, anatomical location, and cellular underpinnings. Specifically, we hypothesized that (i) IBAP extent depends on GA, with earlier birth associated with larger IBAPs, (ii) anatomical locations of IBAPs remain temporally constant along individual development, and (iii) cellular underpinnings of IBAPs involve glial cells. Regarding developmental plasticity, we additionally expected IBAPs to be modified by the child’s early social environment, supported by previous findings linking social environment to neurocognitive outcomes after preterm birth^51–53^. Finally, inspired by neurocognitive heterogeneity, we proposed that IBAP heterogeneity would underpin the cognitive performance variability among preterm born individuals.

To investigate these hypotheses, we focused on CTh and surface area (SA) as paradigmatic outcomes of brain development. First, we employed normative modeling to describe regional CTh and SA developmental trajectories^2^ and to assess spatial heterogeneity of IBAPs across three cohorts, encompassing preterm and full-term neonates, children aged 10 and 12 years, and adults aged 26 and 38 years (for an overview of the study hypotheses, data, and methods, see Supplementary Fig. S1). Second, we tested the temporal consistency of initial injury using longitudinal IBAPs of preterm children and adults and integrated cell density maps derived from the Allen Human Brain Atlas (AHBA) to trace cellular underpinnings of adult IBAPs back to preterm birth. Third, we examined plasticity of IBAPs in response to the environment by linking adult IBAPs with social environmental features of early development. Finally, we tested whether adult IBAPs were associated with IQ after preterm birth. Taken together, we propose a temporally consistent effect of variable initial injury-induced dysmaturation at the macroscopic (i.e., extent and location) and microscopic (i.e., cellular) scale, which is, however, sensitive to early social-environmental influences, leading to individually heterogeneous abnormality patterns after preterm birth.

### Preterm brain development is heterogeneous

Instead of focusing on average dysmaturation outcomes, we sought to demonstrate spatial heterogeneity in brain development after preterm birth across development by using brain MRI data from three cohorts: (i) neonates scanned at term-equivalent age (92 preterm, 375 full-term) from the developing Human Connectome Project (dHCP), children scanned longitudinally at ages 10 and 12 years (191 preterm, 5,762 full-term) from the Adolescent Brain Cognitive Development Study (ABCD-10 and ABCD-12), and (iii) adults scanned at age 26 years (95 preterm, 107 full-term) with a subgroup scanned again at age 38 years (52 preterm, 53 full-term) from the ongoing Bavarian Longitudinal Study (BLS-26 and BLS-38; Supplementary Table S1).

To replicate previous models of average dysmaturation, we used two-sample t-tests to analyze mean CTh differences between preterm and full-term individuals for each cohort for 34 cortical regions of the Desikan-Killiany parcellation^54^. Consistent with previous findings^32,37,40,55^, regional mean CTh was increased in most areas except for the occipital and inferior temporal lobes in preterm neonates, while it was restrictedly decreased in frontal and temporal regions in children, and widely decreased in lateral associative and primary cortices in preterm adults (Fig. 1a: BLS-26; Supplementary Fig. S2a: other cohorts).

**Figure 1:**
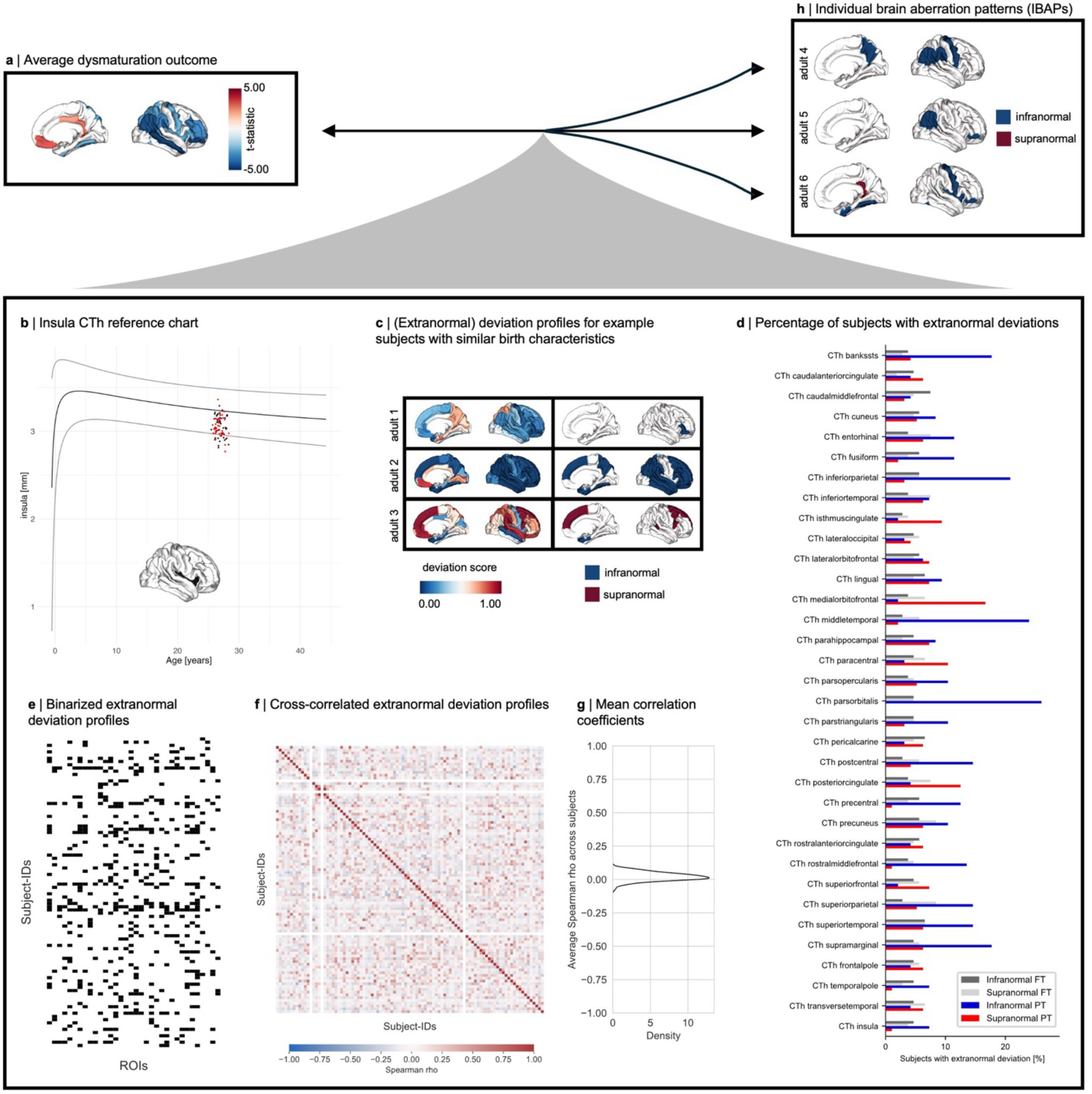
Brain development after preterm birth is individually heterogeneous. **a,** Cortical thickness (CTh) average dysmaturation outcome for 26-year-old adults after preterm birth estimated by two-sample t-tests of between-group differences (p_FDR_ < 0.05) suggests abnormalities shared between preterm subjects. **b,** Individual regional deviations were estimated from a previously published normative framework^2^ and classified as infranormal (i.e., < 5^th^ percentile) or supranormal (i.e., > 95^th^ percentile) for a given cortical region. **c,** Individual deviation score profiles of example subjects with similar gestational age and birth weight. Left column: Deviation scores of all 34 cortical regions, right column: only infra- (i.e., < 5^th^ percentile, dark blue) and supranormal (i.e., > 95^th^ percentile, dark red) deviations. **d,** The percentage of subjects sharing an extranormal deviation in any given region is less than 30 %, illustrating individual heterogeneity of CTh after preterm birth. **e**, Binarized representation of extranormal deviations for each subject. **f,** Spearman correlation matrix of binarized extranormal deviations across subjects. **g,** Distribution of averaged correlation coefficients for each subject with all others.

Next, to refer to spatial heterogeneity of brain aberrations, we analyzed IBAPs for CTh measured as CTh norm deviations based on the BrainChart normative reference framework. This framework has previously been established based on approximately 100,000 subjects to capture typical regional CTh development across the human lifespan^2^. We adopted these reference charts to our datasets by independently estimating random effects of study based on term-born individuals. Normative ranges were operationalized as the range between the 5^th^ and 95^th^ percentiles for a given age and sex. For each individual and each brain region, we calculated deviation scores that quantify how much an individual’s CTh deviates from the normative range (Fig. 1b). A deviation profile refers to the regional distribution of deviation scores across all 34 investigated brain regions. Adapted normative models accurately captured typical development since more than 85 % of full-term individuals resided within the normative range for any given region. To examine whether deviation profiles are heterogeneous across preterm individuals, we first focused on the BLS-26 cohort. Visual comparison demonstrated that the severity and pattern of deviations varied substantially, not only with respect to average dysmaturation outcome (Fig. 1a) but also between individuals, even among those born with similar GA and birth weight (BW; Fig. 1c). Although the majority of individuals (84 % of full-term and 97 % of preterm subjects) showed at least one extranormal deviation, the locations of these deviations were markedly heterogeneous among individuals. No more than 27 % of preterm adults shared extranormal deviations in the same cortical region (Fig. 1d). This result was independent from the normative model used to estimate regional CTh development in the population (Supplementary Methods and Supplementary Fig. S3)^2,56^, suggesting both the robustness of our finding regarding the applied model and distinct patterns of cortical dysmaturation after preterm birth. To investigate potential spatial patterns of extranormal deviations in more detail, we created binarized deviation profiles (i.e., infra- and supranormal deviations are designated with 1, all others with 0; Fig. 1e) and cross-correlated them across preterm subjects to estimate deviation profile similarity (Fig. 1f). Although extranormal deviation profiles show similarities between some preterm subjects (Spearman rho > |0.4|), the average correlation coefficient between one individual’s pattern and all others was very weak (ranging from -0.07 to 0.09), suggesting individually heterogeneous patterns of CTh deviations across cortical regions.

To investigate whether this heterogeneity is observable across development, we analyzed deviation profiles in neonates, children, and adults aged 38 years (Supplementary Fig. S2b-c). The overall prevalence of at least one extranormal deviation varied with age from only less than 65 % in preterm neonates to over 85 % of preterm children and adults, suggesting developmental and/or environmental influences on brain structure. Location patterns remained heterogeneous across all developmental stages, with no more than 27 % of preterm individuals sharing deviations in any given region, suggesting that CTh is heterogeneously altered across development after preterm birth.

To determine whether individual heterogeneity of structural abnormalities after preterm birth also applies to other brain features, we examined regional SA (Supplementary Fig. S4) as well as cerebral tissue volume measures of white matter volume (WMV), grey matter volume (GMV), and subcortical grey matter volume (sGMV; Supplementary Fig. S5). Regional percentile SA trajectories for the entorhinal cortex were not available due to known issues of lower data quality or cortical surface reconstruction for this region^2,57^. Most brain measures showed similar heterogeneity compared to CTh, with only less than 30 % of preterm individuals sharing abnormalities in the same location for regional SA, GMV, and WMV. This suggests that the concept of individual heterogeneity of regional brain measures after preterm birth generally holds for cortical development. In contrast, we observed that up to 40 % of preterm adults deviated in sGMV, which might point to a different mechanism for subcortical development. These findings indicate that brain development after preterm birth is characterized by persistent individual heterogeneity across developmental stages, affecting multiple structural measures of brain development.

### Temporal consistency of preterm dysmaturation

Despite demonstrating that brain development after preterm birth is substantially heterogeneous between individuals, we hypothesized that biological mechanisms of injury-induced dysmaturation underlying these aberrations would be consistent across preterm subjects. Therefore, we examined macroanatomical (i.e., extent and location) and microanatomical (i.e., cellular underpinnings) consistency of deviations across development (Supplementary Fig. S1b, c, and d, respectively).

#### The extent of IBAPs scales with gestational age

To investigate whether the extent of extranormal deviations depends on the severity of prematurity, we first divided the BLS-26 preterm adults into two groups of earlier (GA of ≤ 30 weeks, n = 53) and later birth (GA of > 30 weeks, n = 43). In the earlier born group, up to 36 % of preterm subjects shared a deviation in the same location, while only 19 % of subjects did in the later born group (Supplementary Fig. S6b). Similarly, up to 37 % of preterm neonates born before 30 weeks GA (n = 27) shared the location of supranormal deviations, whereas this was the case for only 18 % of subjects born later in the preterm period (n = 65; Supplementary Fig. S6a). This indicates that earlier birth increases the likelihood of structural alterations. To further support this relationship, we examined whether the number of extranormal brain regions scales with GA. In both preterm neonates (Spearman rho = -0.198, p = 0.059) and adults (Spearman rho = -0.313, p = 0.002), earlier birth was associated with a higher number of extranormal regions (Fig. 2a), demonstrating that the timing of preterm birth influences the extent of extranormal CTh deviations. As a control analysis, we examined two additional measures of the severity of prematurity in the BLS-26 cohort, namely BW and the Duration of Neonatal Treatment Index (DNTI). DNTI provides an estimate for the severity of medical complications after birth. In contrast to BW (Spearman rho = -0.162, p = 0.116), DNTI showed a significant association with the number of infranormal regions (Spearman rho = 0.242, p = 0.018; Supplementary Fig. S6c). These results demonstrate that GA has a substantial impact on the extent of CTh deviations, with earlier birth leading to more widespread patterns of deviations, also later in life.

**Figure 2:**
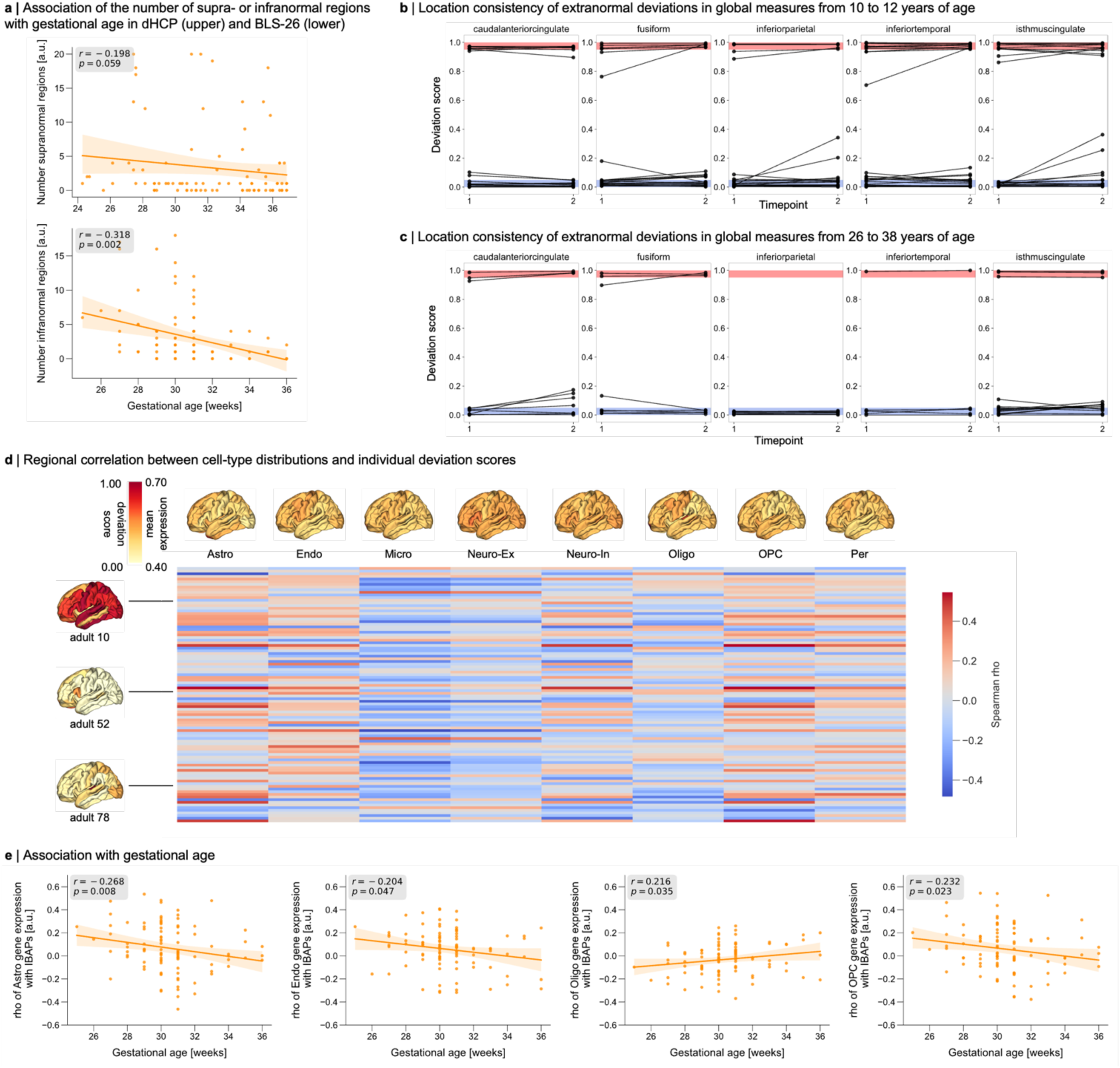
Temporal consistency of IBAP extent, anatomical location, and cellular underpinnings. **a,** The number of supranormal or infranormal CTh deviations per subject was correlated with gestational age for neonate dHCP and adult BLS-26, respectively (dHCP: Spearman rho = -0.198, p = 0.059; BLS-26: Spearman rho = -0.318, p = 0.002). Longitudinal deviation scores of preterm children from ABCD (**b**, i.e., acquired at the ages of 10 and 12 years) as well as preterm adults (**c**, i.e., acquired at the ages of 26 and 38 years) were calculated for regional surface area (SA) in the bilateral Desikan-Killiany parcellation (five example regions are depicted here, a complete representation can be found in Supplementary Fig. S7 - S8). Only deviation scores of subjects that exhibited an extranormal deviation at either timepoint are depicted. Within-subject comparison between the two timepoints demonstrates that the anatomical location of infranormal (blue) and supranormal (red) deviations after preterm birth mostly remain consistent. **d,** Regional Spearman correlation matrix of CTh IBAPs from preterm adults aged 26 years (vertical) with mean expression profile of eight brain cell types (horizontal). **e,** Spearman correlation coefficients for the association between astrocytes, endothelial cells, oligodendrocytes, and OPCs, with individual CTh deviations were significantly related to gestational age in preterm adults (p < 0.05).

#### Constant IBAP locations in individual development

To investigate whether locations of extranormal deviations remain stable over time within the same individual, we compared regional SA deviation scores longitudinally in preterm children (from 10 to 12 years) and in adults (from 26 to 38 years). We focused on SA rather than CTh due to previous studies demonstrating that FreeSurfer estimates of “raw” regional CTh have substantially higher overall variability compared to SA^58^. We observed that preterm subjects who exhibited an extranormal deviation for SA in a particular region at the first timepoint were likely to exhibit a similar deviation in that same region at the second timepoint (Fig. 2b-c for selected regions of ABCD and BLS due to space limitations, Supplementary Fig. S7-8 for all cortical regions of both datasets). For infranormal deviations, 77 % of children and 74 % of adults maintained their patterns between scans. Supranormal deviations showed even higher consistency, with 71 % stability in children and 87 % in adults. While some regions showed higher variability, possibly due to measurement stability issues for these regions^58^ or by non-linear developmental trajectories^2,59,60^, individual patterns generally remained stable throughout childhood and adulthood.

#### Glial underpinnings of adult IBAPs

To investigate whether there are consistent cellular underpinnings of IBAPs across development, we assessed whether the locations of some subjects’ deviations follow the regional distribution of the same glial cells shown to be vulnerable to prematurity-induced injury. Given that CTh variability has been demonstrated to be sensitive to various cellular underpinnings^33,34,61–63^, we focused on CTh deviations in the following analyses. First, we estimated the regional abundance of eight different cell types across 34 cortical regions using gene sets from single-cell gene expression studies of the adult human cortex (Methods, for similar methodologies see ^64,65^) and regional gene expression information from the AHBA^66,67^ (Fig. 2d; Methods). The expression level of cell-specific marker genes has previously been established as an indirect indicator of the relative abundance and/or function of this cell type^68,69^. To bridge macro- and microscale by mapping cell-type marker gene expression patterns to regional CTh deviation scores across the cortical sheet, we conducted regional Spearman correlation analysis across the 34 brain regions for each subject. Thereby, significance was assessed using a permutation test that preserves spatial autocorrelation (“spin test”)^70,71^ (Fig. 2d, filtered for significant associations only in Supplementary Fig. S9). We observed a significant association of subjects’ regional CTh deviation patterns to some cellular distributions (p_spin_ < 0.05). Remarkably, across subjects, we found that the strength of the spatial relationship between cellular distributions and deviation profiles varied as a function of GA only for certain cell types (Fig. 2e). Specifically, earlier preterm birth was linked to a stronger association between regional CTh deviations and the regional distributions of astrocytes, endothelial cells, and oligodendrocyte progenitor cells (OPCs), and to a weaker association with mature oligodendrocytes. This suggests that the locations of deviations in earlier prematurity depends largely on the same glial cells that are known to be vulnerable to initial injury. We interpret this result as a glial consistency of cellular underpinnings of IBAPs.

Altogether, these findings point towards lasting, individually heterogeneous brain deviations, consistent in both macroanatomical extent and location as well as microanatomical association through glial cells.

### Impact of early social environment on adult IBAPs

In addition to consistency in injury-induced dysmaturation processes, we also expected a consistent plastic effect of the early environment on individual structural deviations after preterm birth (Supplementary Fig. S1e). Data from the lifelong BLS cohort allowed us to examine whether an individual’s social environment during childhood might contribute to heterogeneous deviation profiles in adulthood. To summarize CTh deviations, we derived a first principal component (PC1) across all 34 regional CTh deviation scores for preterm adults aged 26 years, capturing 37.2 % of the total variance in CTh deviation scores. Loadings for PC1 were positive across all 34 regions (Supplementary Figure S10a), indicating that this component represents a global trend in CTh deviations across the cortex, with lower PC1 scores reflecting lower deviation scores relative to population norms. As representatives of the early developmental environment, we used family socio-economic status (SES) as a general, widely used measure of the economic and sociological environment components of a developing child^72^, and the Parent-Infant-Relationship Index (PIRI) as a measure of parental social environment specifically^73^. SES was assessed neonatally as a composite score of family profession and parental education based on parent interviews and was classified into low, middle, and high^74^. PIRI describes attachment-related parental concerns and the mother’s current and anticipated relationship problems^73^. A sum score from 0 (good parent-infant relationship) to 8 (poor parent-infant relationship) was calculated based on nurse observations neonatally and an interview at five months after birth.

To analyze the impact of the early social environment on CTh heterogeneity after preterm birth, we correlated the respective variable with CTh PC1 scores. CTh PC1 scores and both SES (Spearman rho = -0.217, p = 0.033) and PIRI (Spearman rho = -0.211, p = 0.044) were significantly associated (Fig 3a), indicating that a more favorable social and economic environment links to higher CTh deviations across the cortex. As control measures for PCA-based integrated individual CTh deviations across the cortex, we used the number of infranormal regions as well as the deviation scores of mean CTh across the whole cortex (Supplementary Fig. S11). In preterm adults, we observed that a lower SES was correlated with a lower deviation score of mean CTh (Spearman rho = -0.223, p = 0.029) and a higher number of infranormal regions per subject (Spearman rho = 0.200, p = 0.051), substantiating the association between early social environment and CTh deviations. Based on previous literature suggesting that a lower SES modifies the relationship between preterm birth and adverse neurodevelopmental outcomes^51–53^ in later life, we tested whether a lower SES also adversely modifies the association between preterm birth and adult IBAPs. Moderation analysis revealed that the association between GA and PC1 scores (r = 0.264, p = 0.009; Fig. 4b) was significantly weakened among individuals with middle (p_one-sided_ = 0.001) and low (p_one-sided_ < 0.001) SES, indicating that an adverse early social environment modifies IBAPs after preterm birth.

**Figure 3:**
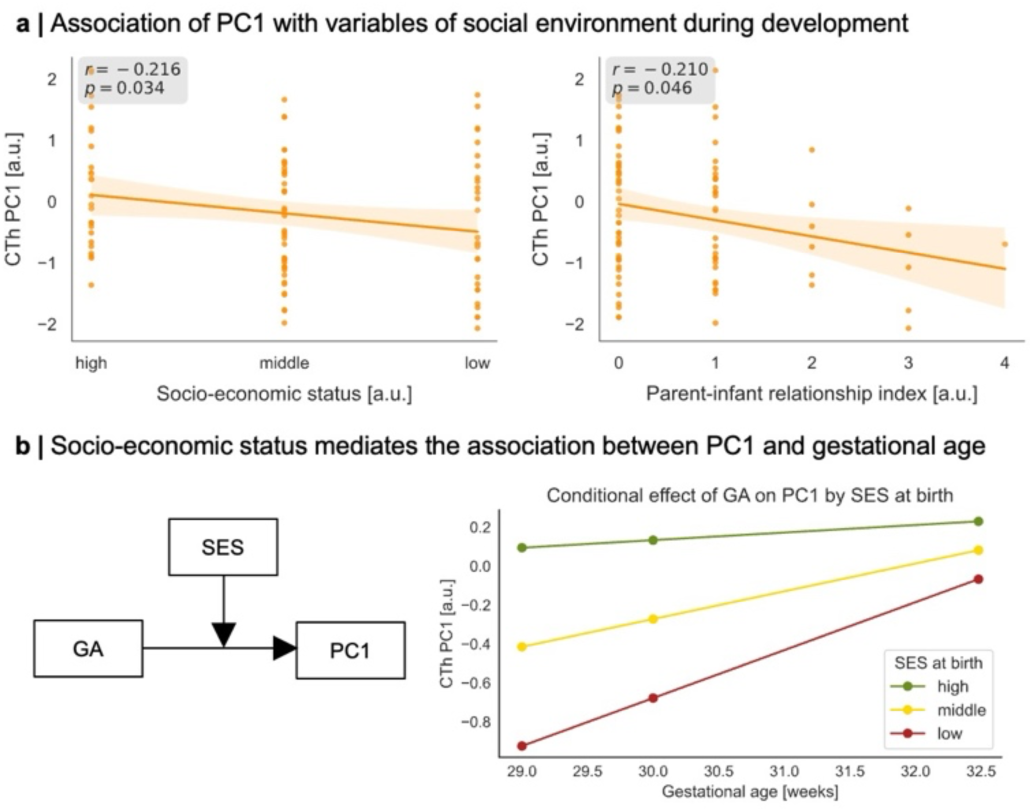
Adult CTh deviations are linked to the social environment during early development. **a,** To capture regional variation of deviation scores across 34 cortical regions, we applied PCA to individual CTh deviations for the BLS-26 dataset (Supplementary Fig. S1e). We correlated the main axis of deviation across regions (i.e., PC1) with early development socio-economic status (SES) and quality of mother-infant relationship (Parent-Infant Relationship Index, PIRI). **b,** The association between GA and CTh PC1 scores were significantly moderated by low (p < 0.001) and middle (p = 0.001) socio-economic status.

**Figure 4:**
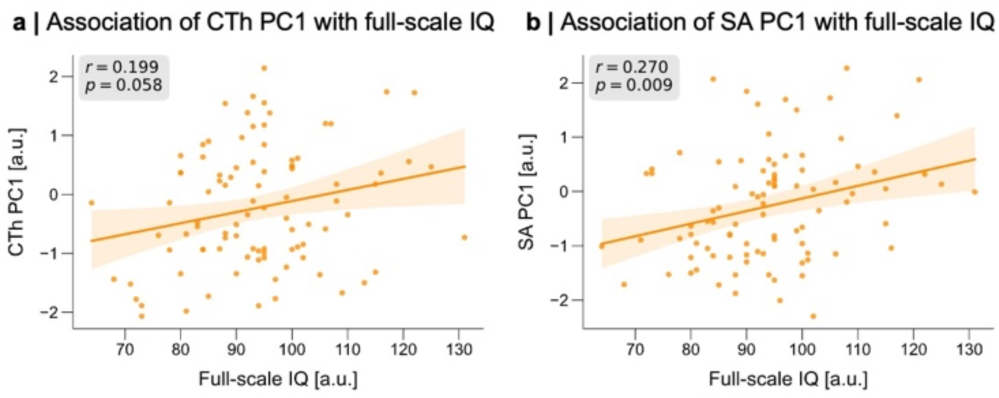
Individual brain deviations after preterm birth are associated with cognitive outcome variability in adulthood. **a**, Spearman correlations between CTh PC1 scores and full-scale IQ in preterm adults aged 26 years. **b**, Spearman correlations between SA PC1 scores and full-scale IQ in preterm adults.

### Impacts on cognitive outcome variability

Finally, to test whether heterogeneity in IBAPs underlies cognitive performance variability among preterm individuals, we associated CTh and SA PC1 scores with full-scale IQ in the BLS-26 cohort. Due to the link of a more favorable social environment with higher CTh PC1 scores, we expected lower PC1 scores to be associated with lower individual IQ in preterm adults. We observed a significant correlation between full-scale IQ and CTh PC1 (Spearman rho = 0.198, p = 0.030). The control analyses associating the number of infranormal CTh regions with IQ confirmed this relationship (Spearman rho = -0.227, p = 0.030; Fig. 4a). Likewise, SA PC1 scores was significantly linked to IQ in preterm adults (Spearman rho = 0.270, p = 0.009; Fig. 4b). This relationship was specific to preterm individuals, with full-term individuals not showing a significant association (CTh: Spearman rho = 0.030, p = 0.760; SA: Spearman rho = 0.041, p = 0.679). Control analyses of correlations between IQ with the deviation score of mean CTh (Spearman rho = 0.194, p = 0.063) as well as with the number of infranormal regions of CTh (Spearman rho = -0.227, p = 0.030) and SA (Spearman rho = -0.214, p = 0.041) reinforced the relationship between a more severely affected cortex with lower IQ (Supplementary Fig. S12). In brief, results indicate that individual structural deviations contribute to the variability of general cognitive performance in adulthood after preterm birth.

## Discussion

This study provides evidence for lasting and heterogeneous brain aberrations after preterm birth, shaped by consistent effects of variable initial injury-induced dysmaturation at macroscopic (i.e., extent and location) and microscopic (i.e., cellular) scales, and influenced by the early social environment. This individual heterogeneity of cortical development, combined with biological mechanistic consistency for injuries and plasticity, integrates two opposing suggestions on preterm brain development: highly heterogeneous neurodevelopment due to varying neurocognitive outcomes versus regular aberrant neurodevelopment based on average dysmaturation outcomes of brain imaging studies. Thus, the concept of broadly altered brain development with widespread, persistent, average-based brain aberrations after preterm birth needs to incorporate the substantial component of individual heterogeneity following temporally consistent but environmentally sensitive trajectories of dysmaturation (Fig. 5).

**Figure 5:**
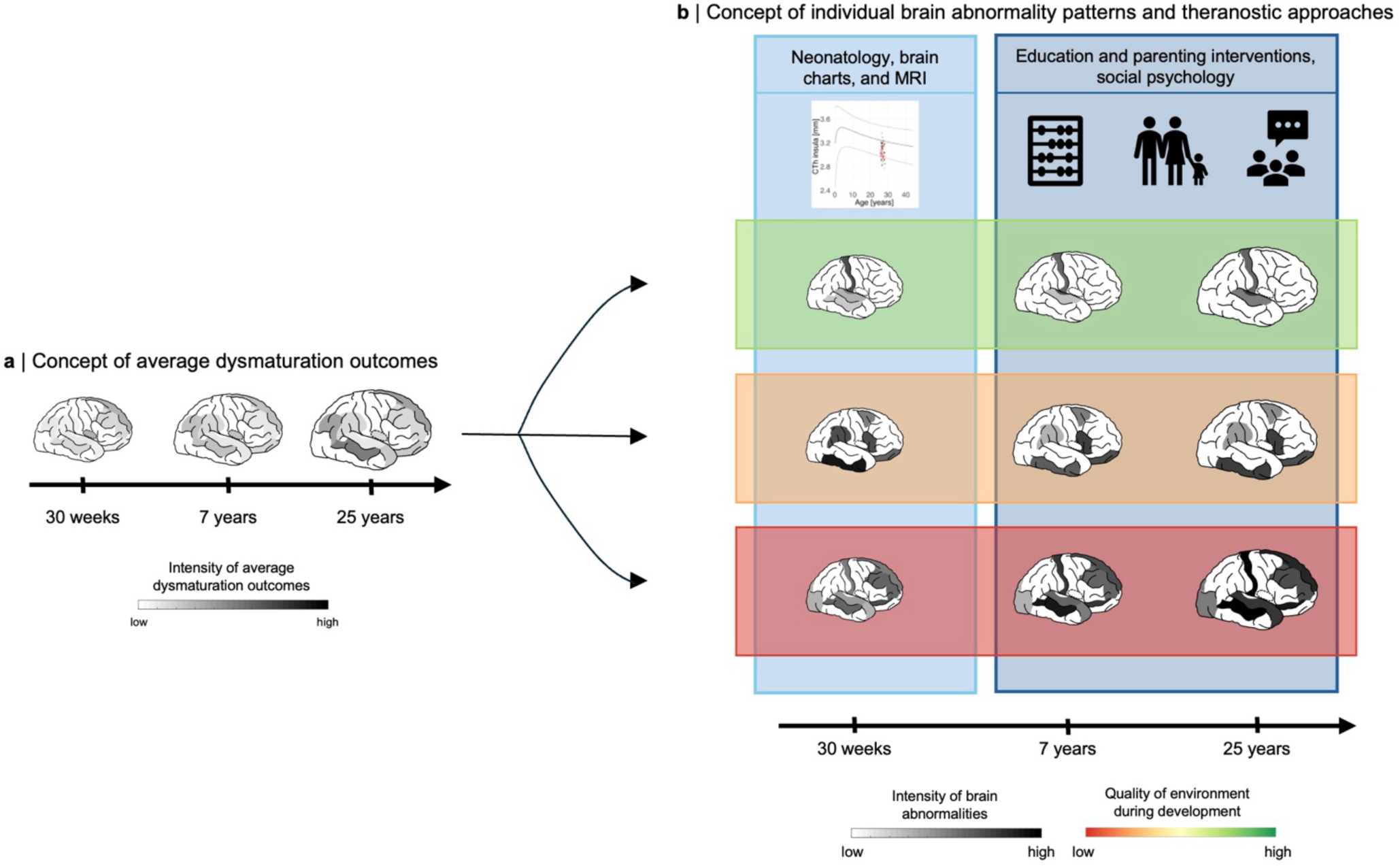
Conceptual transition from average dysmaturation outcome-based view to individual brain abnormality pattern centered model of prematurity. Schematic representation of the two contrasted concepts. **a,** The prevailing view of brain aberrations based on group-level brain imaging studies after preterm birth suggests injury-induced average dysmaturation outcomes, disregarding individual variability. **b,** The extended concept presented here emphasizes the heterogeneity of brain abnormality patterns among preterm individuals, as a result of individual initial injury patterns (indicated by varying extent and location of abnormalities across three arbitrary subject cases), subsequent dysmaturation (indicated by a brain trajectory for each case), and plastic individual changes due to social environment influences (indicated by horizontal boxes around distinctively developing brain trajectories). Critically, the model of heterogeneous and plastic IBAPs suggests specific theranostic approaches on prematurity (blue shaded vertical boxes): on the one hand, neonatology and chart-based brain imaging approaches to identify at-risk infants, and on the other hand, social psychology approaches to social environment to modify developmental trajectories. Thus, heterogenous IBAPs seem to have the potential to frame and integrate approaches of basic neuroscience, neonatology, brain imaging, and social psychology on human prematurity.

Leveraging normative modeling approaches, we identified distinct individual brain abnormality patterns (IBAPs) in preterm individuals across life stages, varying in extent and spatial distribution. While group-level analyses suggest spatially consistent alterations across subjects, individual effects vary substantially (Fig. 1a, Supplementary Fig. S2, S4, S5). Only about a quarter of preterm subjects exhibited an overlapping deviation in the same cortical region, indicating that the degree and location of brain abnormalities in an individual cannot be inferred from average dysmaturation outcomes derived from group-level analyses. This heterogeneity persisted across all examined cohorts, extended beyond CTh to other structural measures, and was robust across different normative reference models (Supplementary Fig. S3), suggesting individual developmental trajectories of the brain after preterm birth. These findings substantially advance the concept of individually heterogeneous brain development after preterm birth, previously proposed by postmortem and imaging studies, which presented high spatial variability of glial cell-mediated white matter lesions between subjects^75–77^. Consistent with our results, previous normative modeling studies have also presented substantial heterogeneity of brain measures in preterm neonates, including CTh and SA^32,49,50^, but without investigating possible manifestations in childhood or adulthood. Our results fill this critical gap by presenting heterogeneous structural aberrations in preterm neonates, children, and adults, fundamentally extending our understanding of brain development after preterm birth.

Despite the considerable heterogeneity of brain aberrations, injury mechanisms regarding the extent, location, and cellular underpinnings of individual abnormality patterns seem to share commonalities (Fig. 2, Supplementary Fig. S6 – S9). Earlier gestational age was correlated with more extranormal deviations in both neonates and adults, suggesting a consistent extent of deviations from birth into adulthood. Considering that previous studies have shown that earlier born preterm individuals are more likely to have brain injuries, extreme individual brain aberrations, and subsequent neurodevelopmental impairments^32,78–80^, these findings are not surprising. Moreover, structural deviations outside the normative range demonstrated remarkable microanatomical stability throughout development, with some regional exceptions potentially due to signal quality issues ^57^ or non-linear maturation trajectories^2,59,60^. Studies that analyzed group average longitudinal trajectories of raw CTh and SA from childhood to early teenage years^38^ and from adolescence to young adulthood^81^ after preterm birth also reported a consistent aberration pattern across individuals in later life. In conclusion, individual cortical aberrations are dependent on gestational age and seem to persist at the same locations into middle adulthood.

Investigating potential cellular-molecular underpinnings, we found that the cell types mediating initial injury effects of preterm birth on individual CTh aberrations are similar across the first half of the lifespan, i.e., involve certain glial cell types. As expected, individually heterogeneous regional CTh deviation patterns were significantly linked to several different cell type profiles, again stressing that CTh aberrations are different across preterm individuals. Critically, however, adults born earlier in the preterm period exhibit deviation patterns more closely aligned with the regional distributions of astrocytes, endothelial cells, and OPCs and less aligned with those of oligodendrocytes. This suggests consistent cellular underpinnings between neonates and adults, at least for adults born earlier in the preterm period. Particularly, glial cells prone to hypoxia-ischemia injury may be damaged around time of birth^11–17,82–87^, with extranormal deviations still being present at the locations where most astrocytes, endothelial cells, and OPCs reside in adulthood. OPCs are highly vulnerable to hypoxia-ischemia and are replenished after damage but fail to differentiate to mature oligodendrocytes in the following, resulting in hypomyelination^12^. This could potentially explain the overlap of IBAPs with OPC but the anti-correlation with oligodendrocyte cell distributions. As cortical thinning is largely dependent on intracortical myelination^33,34^, areas with higher OPC abundance might be more prone to extreme cortical thinning after preterm birth. This interpretation assumes that regional cellular abundance in the adult and infant brain overlap. Yet, previous evidence suggests that transcriptional profiling of cells alters across development, but the regions where cells reside once at place remain mainly conserved after birth^88,89^, substantiating our interpretation. Collectively, these findings reveal that the individual heterogeneity in regional CTh observed in adults born earlier in the preterm period is significantly associated with the regional distributions of glial cells implicated in neonatal brain injury.

Furthermore, we present evidence that IBAPs after preterm birth are modifiable by the early social environment and relevant for cognitive outcome variability, demonstrating an interplay between environment, brain development, and eventually long-term cognitive outcomes. By applying PCA, we captured global CTh deviation as PC1. Individual adult CTh PC1 scores were significantly influenced by a subject’s social environment during infancy (Fig. 3), with medium or low SES moderating the association between GA and CTh PC1 scores. Importantly, this suggests that a more favorable early social environment could mitigate the adverse effects of preterm birth on the brain. As social influences continue throughout childhood including parenting, peer and neighborhood influences^90,91^, further investigation of childhood social influences is warranted. Taken together, we show that individual heterogeneous brain aberrations after preterm birth are not only modulated by common perinatal and social factors, but also underpin cognitive outcome variability.

Overall, our findings suggest that brain development following preterm birth is both highly heterogeneous and temporally consistent, yet remains considerably plastic, which opens important theranostic implications (Fig. 5). As changes in neonatal care have increased survival rates but not cognitive, mental health, or health related quality of life^92–95^, interventions post-discharge across childhood may alter brain variability and cognitive development^96^. On the one hand, our findings stress the necessity of individualized diagnostics, for example using normative approaches of brain imaging measures to better identify at-risk infants. On the one hand, in addition to pharmacological interventions for neonates, comprehensive therapeutic strategies need to incorporate social and developmental interventions that focus on the social and learning environment such as parenting interventions and improved support in early education and schooling.

Clearly, the present work should be considered alongside some important methodological limitations. First, for preterm neonates, we rely on cross-sectional data only. Future studies would benefit from longitudinal data, both for reference chart estimations^97^ and for assessing individual preterm brain developmental trajectories. Second, robust models for early age ranges between birth and 2 years of age are lacking apart from the BrainChart reference charts^2^, hindering validation of neonatal results. Since substantial heterogeneity of regional CTh in preterm neonates has been shown previously^32^, however, there is further evidence based on different methodologies for individual heterogeneity of structural brain aberrations after preterm birth also in infancy. Third, investigation of cellular underpinnings relies on the assumption that the spatial organization of adult neurobiology is reflected in CTh estimates and conserved between humans. Although several pieces of evidence support this notion^61,62,66^, non-invasive tools to investigate this assumption are lacking. Furthermore, the relationship between cell-type specific marker-gene expression and cell abundance varies for different cell types^98^, and cell types are actually more diverse than simplified here^99^.

## Conclusion

In summary, we demonstrate that lasting and widespread altered developmental brain aberrations after preterm birth are individually heterogeneous in their extent and distribution across the brain. Still, the underlying initial injury mechanisms with respect to the extent, location, and cellular underpinnings of individual brain abnormalities patterns seem to be related to gestational age and are temporally consistent from birth until adulthood. Importantly, these abnormalities are modifiable by the early social environment, opening theranostic opportunities. Our work uncovers a new integrative perspective on brain alterations after preterm birth by complementing broadly and widespread altered brain development after preterm birth with highly individual heterogeneous brain alterations.

## Data availability

Neonatal imaging data were collected for the developing Human Connectome Project and are available under restricted access for data privacy reasons and by regulations of the original investigators. Access can be obtained by application to the NIHM Data Archive (https://nda.nih.gov/edit_collection.html?id=3955). Further information on how to gain access can be found on the dHCP study website (https://biomedia.github.io/dHCP-release-notes/). Children imaging data were collected for the Adolescent Brain Cognitive Development Study and are also available under restricted access or data privacy reasons and by regulations of the original investigators. Access can be obtained by application to the NIHM Data Archive (https://nda.nih.gov/abcd/request-access). Adult imaging data were collected in-house and are not publicly available due to ethical considerations but are available from the corresponding author on reasonable request. Normative reference charts for regional CTh and SA from the BrainChart project^1^ will be released upon publication (https://github.com/brainchart/Lifespan). Charts for cerebral volume measures are already publicly available (https://brainchart.shinyapps.io/brainchart/).

## Code availability

All code used to perform the analyses presented here can be found at https://github.com/Melissa1909/preterm-brain-heterogeneity. Code to re-fit pretrained normative BrainChart models will be available upon publication at https://github.com/brainchart/Lifespan.

## Methods

### Ethics

Specific approval for collection and sharing of all data used in this study (brain atlases, BrainChart model, Allen Human Brain Atlas, dHCP, ABCD, and BLS) were provided by local ethics committees; detailed information is available in the cited sources. Informed consent was obtained from each participant and their parents in the case of underage subjects.

### Software

If not stated otherwise, analyses were carried out in a Python 3.11.9 environment.

### Neonatal brain measure estimation from dHCP data

Infant brain measures were obtained from the developing Human Connectome Project^2^ (approved by the National Research Ethics Committee; REC: 14/Lo/1169; 2^nd^ data release). Prior to imaging, informed written parental consent was obtained. Term-born and preterm-born babies were scanned at term-equivalent age (37 – 45 weeks postmenstrual age) during natural unsedated sleep at the Evelina London Children’s Hospital. T2-weighted (T2w) scans with the following parameters were acquired: TR = 12 s, TE = 156 ms, SENSE = 2.11/2.58 (both axial/sagittal) with in-plane resolution of 0.8 × 0.8 mm (1.6 mm slice thickness, 0.8 mm overlap). A 3D motion corrected sensitivity encoding reconstruction was applied, resulting in 0.5 mm isotropic resolution images^3^. Infant preparation and imaging have been previously described in detail^2^. Term-born subjects were excluded if admitted to neonatal intensive care unit or if scans revealed a significant intracranial abnormality (e.g., acute infarction and parenchymal hemorrhage but not punctate white matter lesions, small subependymal cysts/hemorrhages in the caudothalamic notch, mildly prominent ventricles, or widening of the extra-axial CSF). Preterm subjects were only excluded if they had major congenital malformations or diagnosed genetic disorders. Supplementary Table 1 shows demographic data of the final study sample of 375 term-born and 92 preterm-born individuals. Thickness and surface area maps were retrieved from automatically processed T2w images^4^ and were parcellated into 34 bilateral cortical regions using the Desikan-Killiany-compatible M-CRIB-S(DK) atlas^5,6^ and averaged across hemispheres.

### Child brain measure estimation from ABCD data

Brain measures for children were obtained directly from 4^th^ release of the Adolescent Brain Cognitive Development (ABCD) Study® (https://doi.org/10.15154/1523041)^7^. T1-weighted (T1w) and T2-weighted (T2w) MRI scans were acquired at different 3T scanners at 21 sites, with the parameters that are listed on the study website (https://abcdstudy.org/images/Protocol_Imaging_Sequences.pdf). MRI data from baseline (acquired at around age 10 years) and the 2-year-follow-up (acquired at around age 12 years) were preprocessed using FreeSurfer 7.1.1 by the ABCD study team. Details about image acquisition and processing can be found in the original manuscript by Casey et al.^7^ and in the release manuscript (https://doi.org/10.15154/1523041). Following quality control procedures previously established in normative modeling studies^1,8^, we used FreeSurfer’s Euler Index (EI), an automated, quantitative measure of cortical reconstruction quality^9,10^, to estimate surface defects in cortical surface reconstructions. Following Lotter et al.^11^, we excluded subjects that had at least one missing value for the investigated brain measure (i.e., CTh or SA), exceeded a threshold of Q3+IQRx1.5 calculated in each sample across timepoints^9,12^, or failed the manual quality ratings provided in the ABCD dataset. Gestational age was derived from a parent questionnaire, asking “About how many weeks premature was the child when they were born?” (included in file *abcd_devhxss01.txt* of the 4^th^ data release). We determined gestational age as 40 weeks minus the number of weeks the baby was said to be born premature. Since this estimation is rather vague, we only included subjects born more than 7 weeks premature (i.e., gestational age ≤ 32 weeks) as preterm and born at or after week 37 as full-term. Moreover, only subjects with longitudinal assessments were included. Regional CTh estimates were extracted for 191 preterm and 5,762 full-term participants (Supplementary Table S1), regional SA estimates for 148 preterm and 4,929 full-term participants (i.e., due to a higher number of missing values for SA). Estimates were averaged across hemispheres.

### Adult brain measure estimation from BLS data

The Bavarian Longitudinal Study (BLS) is a geographically defined, whole-population sample of individuals born very preterm (VP; < 32 weeks of gestation) and/or with very low birth weight (VLBW; < 1,500 g) and full-term (FT; > 37 weeks of gestation) controls that were followed from birth between January 1985 and March 1986 until adulthood^13,14^. 682 infants were born VP and/or with VLBW. Informed written consent from a parent and/or legal guardian was obtained. From the initial 916 FT born infants born at the same obstetric hospitals who were alive at 6 years, 350 were randomly selected as control subjects within the stratification variables of sex and family socioeconomic status in order to be comparable with the VP/VLBW sample. Of these, 411 VP/VLBW individuals and 308 controls were eligible for the 26-year follow-up assessment. 260 of the VP/VLBW group and 229 controls participated in psychological assessments^15^. All subjects were screened for MR-related exclusion criteria, including (self-reported): claustrophobia, inability to lie still for > 30 minutes, unstable medical conditions (e.g., severe asthma), epilepsy, tinnitus, pregnancy, non-removable MRI-incompatible metal implants and a history of severe CNS trauma or disease that would impair further analysis of the data. However, the most frequent reason not to perform the MRI exam was that subjects declined to participate. Finally, 101 VP/VLBW subjects and 111 FT controls underwent MRI at 26 years of age. The MRI examinations took place at two sites: The Department of Neuroradiology, Klinikum rechts der Isar, Technische Universität München, (n = 145) and the Department of Radiology, University Hospital of Bonn (n = 67). The study was carried out in accordance with the Declaration of Helsinki and was approved by the local ethics committee of the Klinikum rechts der Isar, Technische Universität München and the University Hospital Bonn. All study participants gave written informed consent. At both sites, MRI data acquisition was performed on Philips Achieva 3 T TX systems or Philips Ingenia 3 T systems using an 8-channel SENSE head coil. A high-resolution T1w 3D-MPRAGE sequence (TI = 1300 ms, TR = 7.7 ms, TE = 3.9 ms, flip angle = 15°, field of view = 256 mm × 256 mm, reconstruction matrix = 256 × 256, reconstructed isotropic voxel size = 1 mm^3^) was acquired. Further details about the dataset are described elsewhere^16–19^. For a subset of the participants, a second T1w image was acquired at the age of 38 on a Philips Ingenia Elition X 3 T scanner at the Klinikum rechts der Isar, Technical University of Munich. A 3D-MPRAGE sequence with the following parameters was acquired: TI = 692 ms, TR = 8.7 ms, TE = 3.8 ms, flip angle = 8°, field of view = 256 mm × 240mm x 161 mm, reconstruction matrix = 256 × 256, compressed-SENSE factor = 3, reconstructed isotropic voxel size = 1 mm^3^. For both acquisition timepoints, surface-based parcellation to extract regional CTh and SA in the Desikan-Killiany atlas^6^ was conducted using FreeSurfer (version 7.3.2; http://surfer.nmr.mgh.harvard.edu/). Global measures of cerebrum tissue volume were also retrieved from FreeSurfer. For quality control, FreeSurfer’s EI was extracted. Scans that had EI greater than 4 median absolute deviations above the median EI (n = 8) were excluded. Demographics of the final adult study samples are summarized in Supplementary Table 1. Regional CTh and SA were averaged across hemispheres to match the normative modeling framework of BrainChart, resulting in 34 bilateral cortical ROIs. GA and BW were obtained from obstetric records.

### Data harmonization

As the ABCD-10, ABCD-12, and BLS-26 datasets were acquired on different scanners, bilateral regional CTh and SA as well as GMV, sGMV, and WMV values were retrospectively harmonized to remove variance related to different hardware and scanner sites. Since the scanners used for data acquisition of BLS-26 and BLS-38 were differing and BLS-38 was acquired with a single scanner, only the cross-sectional data of BLS-26 were harmonized with NeuroCombat (version 0.2.12 in Python; https://github.com/Jfortin1/neuroCombat)^20^. NeuroCombat has been repeatedly shown to successfully remove scanner variation in diffusion tensor imaging^21^, structural^20^, and functional^22^ MRI data while preserving biological variance. In this algorithm, an empirical Bayesian framework is applied to estimate additive and multiplicative scanner site effects using parametric empirical priors. To adjust for both effects, the data is then subtracted by the additive and divided by the multiplicative effect parameters^20,23^. In our application, we preserved biological variation attributed to sex, prematurity, and age while controlling for the effect of four scanners used for data acquisition. For ABCD-10 and ABCD-12, an extension of NeuroCombat for longitudinal brain imaging data, longCombat (version 0.0.0.9 in R 4.4.1), was used, which has been shown to be more powerful in detecting longitudinal change than cross-sectional NeuroCombat^24^. One of 21 sites stopped data acquisition after baseline and was thus not included in our dataset. We adjusted site effects for 20 sites, while preserving biological variation attributed to sex, prematurity, and age. The application of longCombat requires that the hardware used is the same at both acquisition times, which was not the case in BLS. The dHCP and BLS-38 years datasets were not harmonized for scanner effects since these data were acquired at the same scanner.

### Assessment of group average differences between preterm and term-born subjects (**Fig. 1a**)

To estimate average dysmaturation outcomes for bilateral regional CTh and SA, a linear regression model correcting for age and sex was fit for each cohort. Results were adjusted for the False Discovery Rate (FDR)^25^.

### Obtaining individual brain abnormality patterns (IBAPs; Fig. 1b)

Normative reference charts for bilateral regional CTh, SA, as well as for global measures (i.e., GMV, sGMV, and WMV) were obtained from the BrainChart project^1^, in which GAMLSS-based normative models were computed over the human lifespan as a function of age and sex from about 100,000 MRI scans. To adapt the reference charts to the datasets used for this study, random effects of study were calculated based on full-term individuals. Normative ranges were operationalized as the range between the 5^th^ and 95^th^ percentile. Based on the adapted normative models for each brain measure, a deviation score between 0 and 1 was then calculated for each individual and each cortical region. This deviation score represents the percentile of this person’s CTh or SA value for the respective cortical region in relation to the normative range. Subjects falling outside the limits defined as the normal range were classified as infranormal (i.e., below the 5^th^ percentile) or supranormal (i.e., above the 95^th^ percentile) for a given region.

### Number of extranormal deviations (Fig. 1d)

To assess the number of subjects per group overlapping in their extreme deviation per region, the percentage of subjects within a group exhibiting an extranormal (i.e., infranormal or supranormal) deviation in a single region was calculated for each cohort.

### Assessing spatial heterogeneity of IBAPs after preterm birth (Fig. 1e-g)

To evaluate whether brain aberrations after preterm birth are spatially heterogeneous, we binarized IBAPs of preterm subjects for each cohort, so that deviations below the 5^th^ or above the 95^th^ percentile for any given region were designated with 1 and all others with 0. Next, Spearman correlation was used to cross-correlate binarized IBAPs across subjects. To estimate the similarity of a preterm individual with all others, the average correlation coefficient for a subject to all others was computed.

### Extent consistency (Fig. 2a)

To determine whether the number of extranormal deviations depends on gestational age (GA), preterm individuals of the dHCP and BLS-26 cohorts were grouped into earlier birth (i.e., GA of ≤ 30 weeks) or later birth (i.e., GA of > 30 weeks) before calculating the percentage of subjects with extranormal deviations (Fig. 4a). Furthermore, Spearman’s rank correlation was used to associate the number of extranormal regions with gestational age for preterm subjects of the dHCP and BLS-26 cohort (Fig. 4b).

### Anatomical location consistency along development (Fig. 2b-c)

To determine whether anatomical locations of extranormal deviations (i.e., below 5^th^ or above 95^th^ percentile) are temporally constant along development, we calculated deviation scores for the longitudinal subsamples of ABCD (i.e., 191 PT and 5,762 FT), acquired at around ages 10 and 12, and BLS (i.e., 52 PT and 53 FT), acquired at around ages 26 and 38. To focus on clinically relevant deviations, only subjects who showed an extranormal deviation at a minimum of one of the two timepoints were selected. We compared whether extranormal deviations remained extranormal over time.

### Microarray gene expression

Regional microarray expression data were obtained from 6 postmortem brains provided by the AHBA (https://human.brain-map.org/)^26^. The *abagen*toolbox (version 0.1.3; https://github.com/rmarkello/abagen) was used to correctly match AHBA gene expression data with native donor space MRI images following published recommendations^27^ as follows:

- Probe aggregation: Microarray probes were aggregated to genes using data provided by Arnatkevičiutė et al.^27^. Probes not matched to a valid Entrez ID were discarded. When multiple probes indexed the expression of the same gene, we selected the probe with the highest mean correlation across donor pairs (i.e., differential stability^28^)
- Intensity-based filtering: Probes were filtered based on their expression intensity relative to background noise^29^. Probes with intensity less than the background in at least 50 % of the samples across donors were discarded.
- Distance threshold: Samples were assigned to brain regions, using Montreal Neurological Institute (MNI) coordinates generated via nonlinear registrations (https://github.com/chrisfilo/alleninf), by minimizing the Euclidean distance between the MNI coordinates of each sample and the nearest surface vertex. Samples where the Euclidean distance to the nearest vertex was more than 2 standard deviations above the mean distance for all samples belonging to that donor were excluded. To reduce the potential for misassignment, sample-to-region matching was constrained by hemisphere and gross structural divisions (i.e., cortex, subcortex/brainstem, and cerebellum, such that e.g., a sample in the left cortex could only be assigned to an atlas parcel in the left cortex)^27^. All tissue samples not assigned to a brain region in the provided atlas were discarded.
- Sample normalization: To address inter-subject variation, tissue sample expression values were normalized across genes using a scaled robust sigmoid function^30^:
- Gene normalization: Genes were normalized across tissue samples, again using a scaled robust sigmoid
- Regional gene expression matrix: Samples assigned to the same brain region were averaged separately for each donor and then across donors, yielding a regional expression matrix in the 68-region surface-based Desikan-Killiany parcellation^6^.

### Cell type-specific gene expression maps

As there is currently no available means to measure cellular abundance in vivo, data from multiple single-cell studies of the adult human cortex^31–35^ were compiled previously^36^ to generate a set of cell type-specific gene markers in the adult human brain for the following cell-types: astrocytes, endothelial cells, microglia, oligodendrocyte progenitor cells (OPCs), oligodendrocytes, excitatory neurons, and inhibitory neurons. Cell type-specific maps representing cellular abundance across the cortex were generated by calculating the mean regional expression score for each gene set in the preprocessed AHBA bulk microarray dataset. Expression was averaged across hemispheres to match the bilateral 34 region setting of regional CTh deviation scores.

### Spatial correlation between cell-type gene expression and CTh deviations and relation to gestational age (Fig. 2d-e)

To explore the cellular correlates of individual CTh heterogeneity in preterm 26-year-old adults, we spatially aligned cell type-specific gene expression maps with individual deviation score patterns using spatial Spearman correlation across 34 cortical regions. Whereas correlation coefficients for analyses of two spatial maps are meaningful, parametric p-values are not as they are influenced by the spatial autocorrelation in spatial systems such as the brain, leading to inflated p-values and subsequently to increased familywise error rates across analyses^37^. To circumvent this problem, we applied spatial autocorrelation preserving permutation tests (“spin tests”) to assess the statistical significance of these correlations^37,38^. In brief, random rotations are applied to spherical projections of the brain to calculate a null distribution of correlation coefficients between the two brain maps. The original correlation coefficient is then compared to this null distribution. This method maintains the existing spatial autocorrelation of a brain map by substituting the original values with those from the nearest rotated coordinate^37,38^. The analyses were implemented in Python using the ENIGMA toolbox^39^ (https://github.com/MICA-MNI/ENIGMA). To evaluate the relationship between the spatial covariation of deviation scores and cell type abundance with GA, we used Spearman’s rank correlation of Spearman correlation coefficients between individual deviation scores and cell type-specific gene expression with GA.

### Dimension reduction of individual deviation scores across cortical regions

To capture regional variability of deviation scores across 34 cortical regions, dimension reduction using Principal Component Analysis (PCA) was performed with the *scikit-learn* package (version 1.5.2) in Python. PC1 loadings as well as explained variances for each component were extracted from the calculations.

### Perinatal, social environmental, and behavioral variables for BLS-26

For the BLS-26 cohort, several variables were used to investigate the influences on and consequences of IBAPs in more detail. The duration of neonatal treatment indices (DNTI) was used to quantify the duration of intensive care treatment after birth, providing an estimate of medical complications after birth^40,41^. For this, care level, respiratory support, feeding dependency, and neurological status (mobility, muscle tone, and neurological excitability) were assessed daily and rated on a 4-point scale (0-3)^42^. DNTI is considered the duration of neonatal treatment in days. To estimate the social environment during early development, family socio-economic status (SES) at birth was measured via parental interviews within 10 days after childbirth and computed as a weighted composite score considering the profession of the self-identified head of the family and the highest education held by either parent^43^. Based on this evaluation, SES was classified into high, middle, and low. The Parent-Infant Relationship Index (PIRI) between mother and child was used to quantify attachment-related parental concerns and feelings as well as the parent’s current and anticipated relationship problems^41^. A score ranging from 0 (good) to 8 (poor parent-infant relationship) was calculated based on nurse observations neonatally and an interview at five months after birth. For adults, global cognitive performance at the age of 26 years was assessed with the “Wechsler Intelligenztest für Erwachsene”, the German adaptation of the Wechsler Adult Intelligence Scale, third edition^44^. Results were used to derive full-scale IQ^14,15^.

### Correlations of individual CTh deviations with social environmental and behavioral variables (Fig. 3)

To assess the relations between CTh deviation variability (i.e., PC1) and social environmental or behavioral variables, a Spearman’s rank correlation was performed (*scipy*, version 1.14.1).

### Moderation analysis of SES on the relation between GA and PC1 (Fig. 3b)

To test whether social environment during development mediates the influence of premature birth on the principal component of individual CTh deviations (PC1), a moderation analysis was performed using the PROCESS toolbox (version 4.2)^45^ for R. GA was treated as the independent variable, PC1 as the dependent variable, and SES at birth as the moderator. A test for X by M interaction was performed, and code for visualization was automatically generated. Variables were not mean-centered. One-sided p-values of p < 0.05 were considered significant for the primary interaction effect. Bootstrapping (n = 5,000) was used to estimate 95 % confidence intervals.

### Plots

All plots were generated with Python. Cortical renderings were generated using the ENIGMA toolbox^39^ (https://github.com/MICA-MNI/ENIGMA). Other plots were created using *matplotlib* (version 3.9.2) and *seaborn* (version 0.13.2). Confidence intervals were estimated with bootstrap resampling (n = 10,000). For Fig. 5, brain plots were created with the Simple Brain Plot toolbox (https://github.com/dutchconnectomelab/Simple-Brain-Plot) in MATLAB 2023b (The MathWorks Inc., https://www.mathworks.com).

## Acknowledgements

We thank all current and former members of the Bavarian Longitudinal Study Group who contributed to general study organization, recruitment, data collection, and management as well as subsequent analyses, including (in alphabetical order) Barbara Busch, Stephan Czeschka, Claudia Grünzinger, Paula Rebecca Hippen, Christian Koch, Diana Kurze, Sonja Perk, Andrea Schreier, Antje Strasser, Julia Trummer, and Eva van Rossum. We thank the staff of the Department of Neuroradiology in Munich and the Department of Radiology in Bonn for their help in data collection. Most importantly, we thank all study participants and their families for their efforts to take part in this still ongoing study. Furthermore, we thank Clara Kretschmer for providing help with figure generation using Inkscape.

## Author contributions

M. Thalhammer and C. Sorg designed the study, wrote and edited the manuscript. M. Thalhammer conducted analyses. A. Neubauer contributed to data analysis. J. Seidlitz, M. A. Di Biase and D. Hedderich helped to design the study and provided valuable feedback on data analysis and study design. A. Menegaux, B. Schmitz-Koep, M. Daamen, H. Boecker, D. Wolke, and P. Bartmann were involved in data collection of the adult data set. All other authors made substantial contributions to the conception or design of the work, the acquisition, analysis or interpretation of data, the creation of new software used in the work, or drafted or substantively revised the Article.

