## Supplementary Information for "Heterogeneous, temporally consistent, and plastic brain development after preterm birth"

#### Common nomenclature

| | |
|---|---|
| a.u. | arbitrary unit |
| ABCD | Adolescent Brain Cognitive Development Study |
| AHBA | Allen Human Brain Atlas |
| Astro | Astrocytes |
| BLS | Bavarian Longitudinal Study |
| BW | Birth weight |
| CTh | Cortical thickness |
| CTV | Cerebral tissue volume |
| dHCP | Developing Human Connectome Project |
| DNTI | Duration of neonatological treatment index |
| Endo | Endothelial cells |
| FDR | False discovery rate |
| FT | full-term |
| g | Grams |
| GA | Gestational age |
| GMV | Grey matter volume |
| IBAPs | Individual brain abnormality patterns |
| Micro | Microglia |
| Neuro-Ex | Excitatory neurons |
| Neuro-In | Inhibitory neurons |
| Oligo | Oligodendrocytes |
| OPC | Oligodendrocyte progenitor cells |
| PC1 | Principal component 1 |
| PCA | Principal component analysis |
| Per | Pericytes |
| PIRI | Parent-infant relationship index |
| PT | preterm |
| ROIs | Regions of interest |
| SA | Surface area |
| SES | Socio-economic status |
| sGMV | Subcortical grey matter volume |
| VLBW | Very low body weight (< 1,500 g) |
| VP | Very preterm (< 32 weeks of gestation) |
| WMV | White matter volume |

#### Supplementary methods

##### Deviation score estimation using different normative reference charts by Rutherford et al. for BLS-26

Several research groups have addressed the need for well-defined reference models to quantify the variability of brain measurements across the lifespan[1–3]. To validate our results of individual brain abnormality pattern (IBAP) heterogeneity based on predictions from the BrainChart framework[2] with predictions from another popular, population-based, pretrained model, we deployed the braincharts framework by Rutherford and colleagues[1]. Due to the similar naming of the two frameworks, the BrainChart model by Bethlehem and colleagues[2] used for the main analysis will be termed "Bethlehem-framework", the braincharts model by Rutherford and colleagues[1] used for the control analysis will be termed "Rutherford-framework" in the following. Based on a large neuroimaging cohort of about 46,000 subjects from 59 sites, the Rutherford-framework also provides Desikan-Killiany-based regional CTh and SA estimates for the human lifespan from 2-100 years. Pretrained models were obtained from the developers (https://github.com/predictive-clinical-neuroscience/braincharts) and adapted to the BLS-26 dataset using full-term adults. Subsequently, predictions for the preterm cohorts were obtained, resulting in Z-scores of individual CTh or SA deviations, respectively. Analogue to the main analysis, subjects were classified as (i) infranormal, i.e., < $5^{th}$ percentile corresponding to $Z < -1.645$, (ii) supranormal, i.e., > $95^{th}$ percentile corresponding to $Z > 1.645$, or (iii) normal for each cortical region. For each region, significant extranormal deviations were calculated as described in the Methods section.

Percentages of significant deviations per unilateral cortical region for preterm adults are visualized in Supplementary Fig. S3. In accordance with the main analysis, not more than 27 % of preterm adults significantly deviate from the norm in regional CTh or SA for any one unilateral region, corroborating the notion of substantial heterogeneity among preterm subjects with respect to regional cortical thickness.

### Supplementary figures

#### S1: Study overview

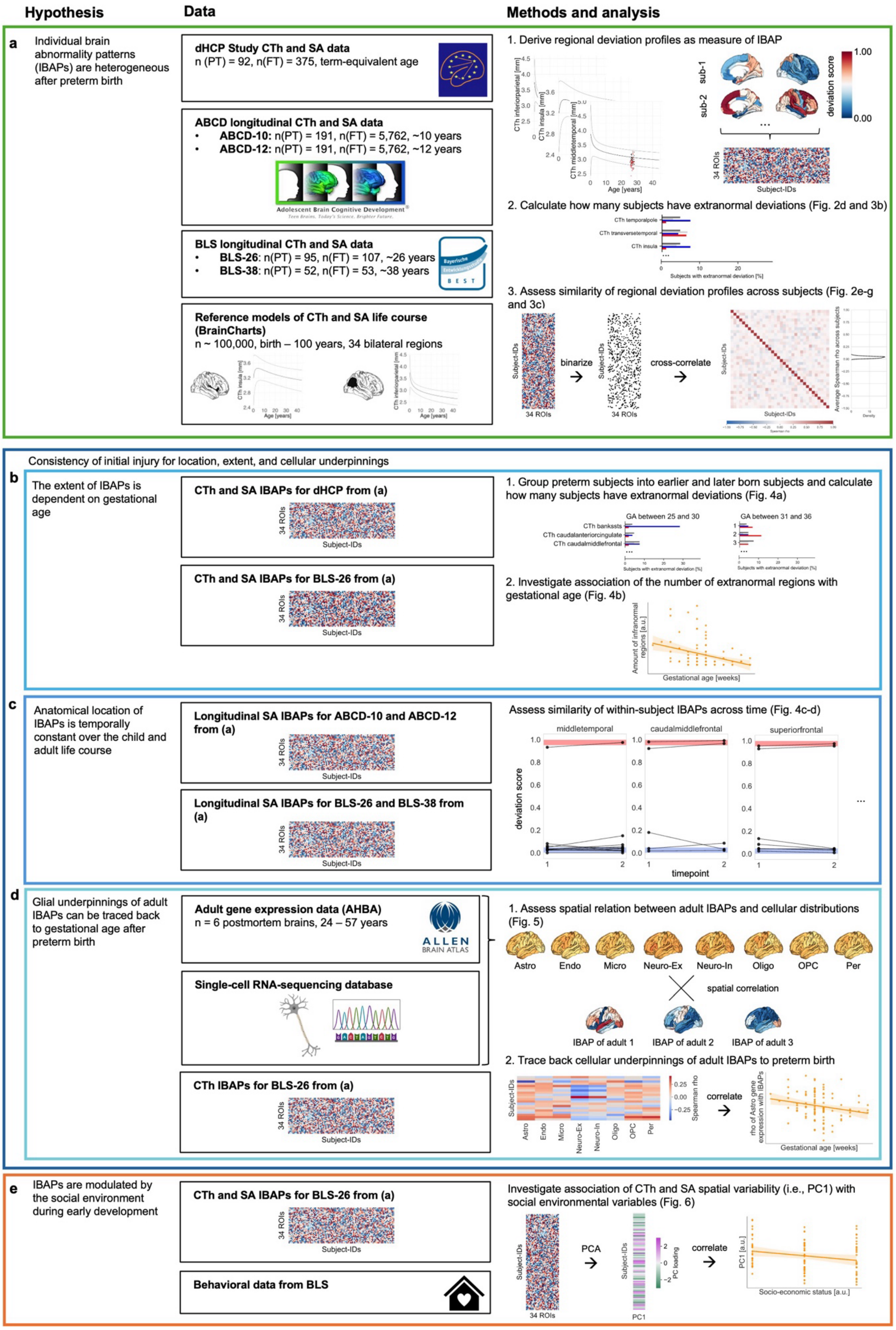

**Supplementary Figure 1: Study overview.** Schematic of the study workflow, from hypotheses (left) to data sources (middle) to analysis steps (right). **a,** Cross-sectional and longitudinal cortical thickness (CTh) and surface area (SA) data were obtained from three developmental cohorts, including preterm and full-term participants. Data were parcellated into 34 bilateral cortical regions. Population CTh and SA life course trajectories were extracted from a normative model. For each participant, individual regional deviations from population life courses were classified as infranormal (i.e., < $5^{th}$ percentile) or supranormal *(i.e.,* > $95^{th}$ percentile) for each cortical region. The regional deviation profile of a certain modality is defined as the measure of the individual brain abnormality pattern (IBAP) for a given participant. (1.) IBAP heterogeneity was assessed in two ways: by quantifying the number of subjects with extranormal deviations in each region (2.), and by measuring spatial similarity of regional deviation profiles across subjects (3.). For the latter, binarized regional deviation profiles were cross-correlated to determine the average correlation between each subject's IBAP with all others. **b,** Consistency in initial brain injury following preterm birth was examined in terms of extent, location, and cellular underpinnings. Extent consistency was examined by relating a subject's number of extra-normal regional deviations to their gestational age. **c,** Location consistency was examined by within-subject longitudinal comparison of IBAPs along childhood and adulthood. **d,** Cellular underpinning consistency was examined by linking gestational age with the spatial correlation of eight brain cell type distributions with adult IBAPs, respectively. **e,** To investigate the developmental plasticity of IBAPs, we linked adult spatial variability of IBAPs to social environmental factors during early life.

#### S2: Individual heterogeneity of regional cortical thickness after preterm birth across cohorts

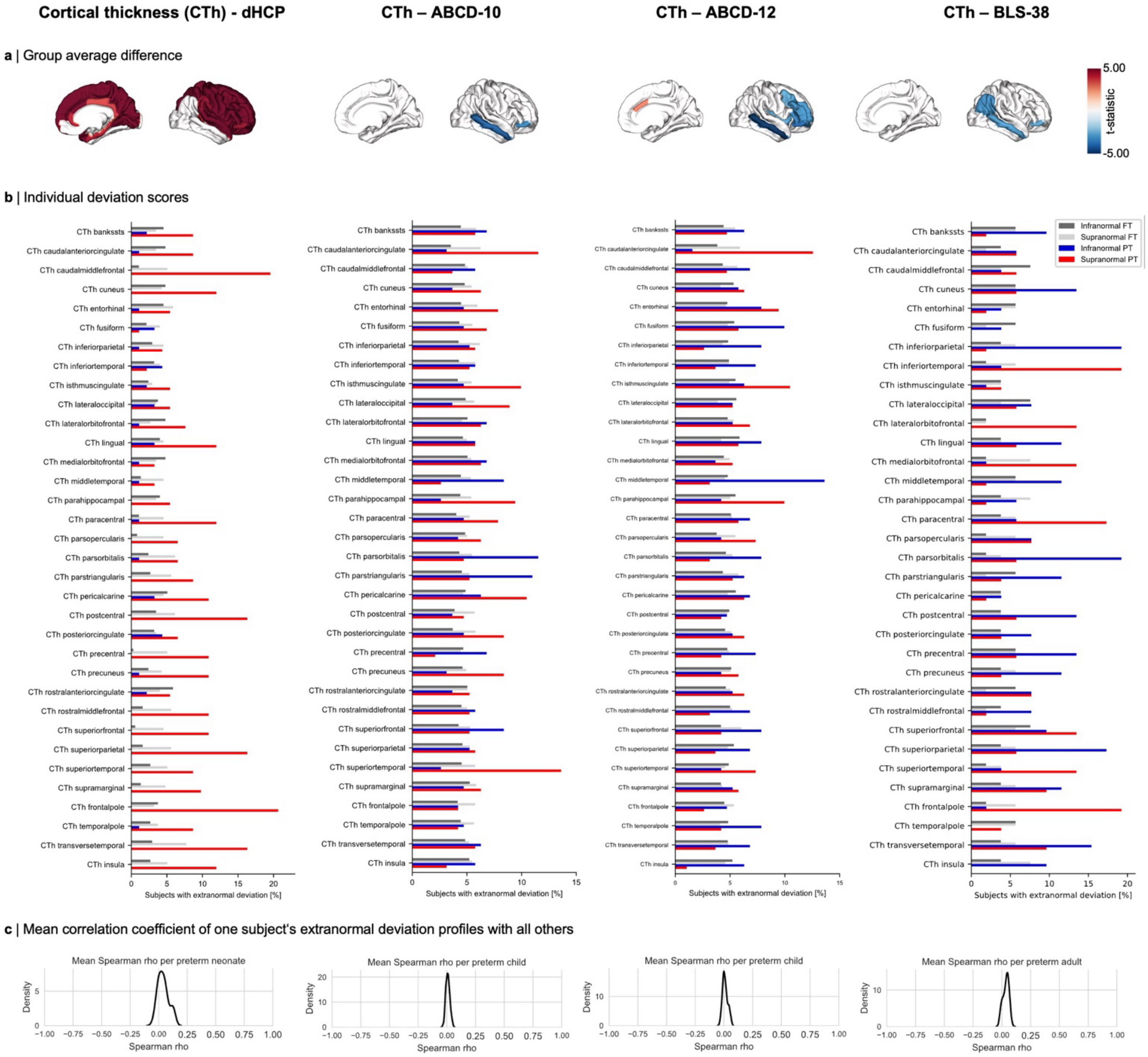

**Supplementary Figure 2: Individual heterogeneity after preterm birth is lasting across age groups. a,** Cortical thickness (CTh) average dysmaturation outcome after preterm birth estimated by T-statistics ($p_{FDR} < 0.05$). **b,** Bar plots showing the percentage of subjects sharing an extranormal deviation in any given region. An overlap of less than 20 % suggests substantial heterogeneity between subjects. **c,** Distribution of averaged correlation coefficients of binarized extranormal deviation profiles for each subject with all others. Results are shown for neonates (dHCP), children (ABCD-10 and ABCD-12), and adults (BLS-38). The shown plot elements correspond partly to Fig. 1.

#### S3: Extranormal individual deviations estimated with a different population reference normative model

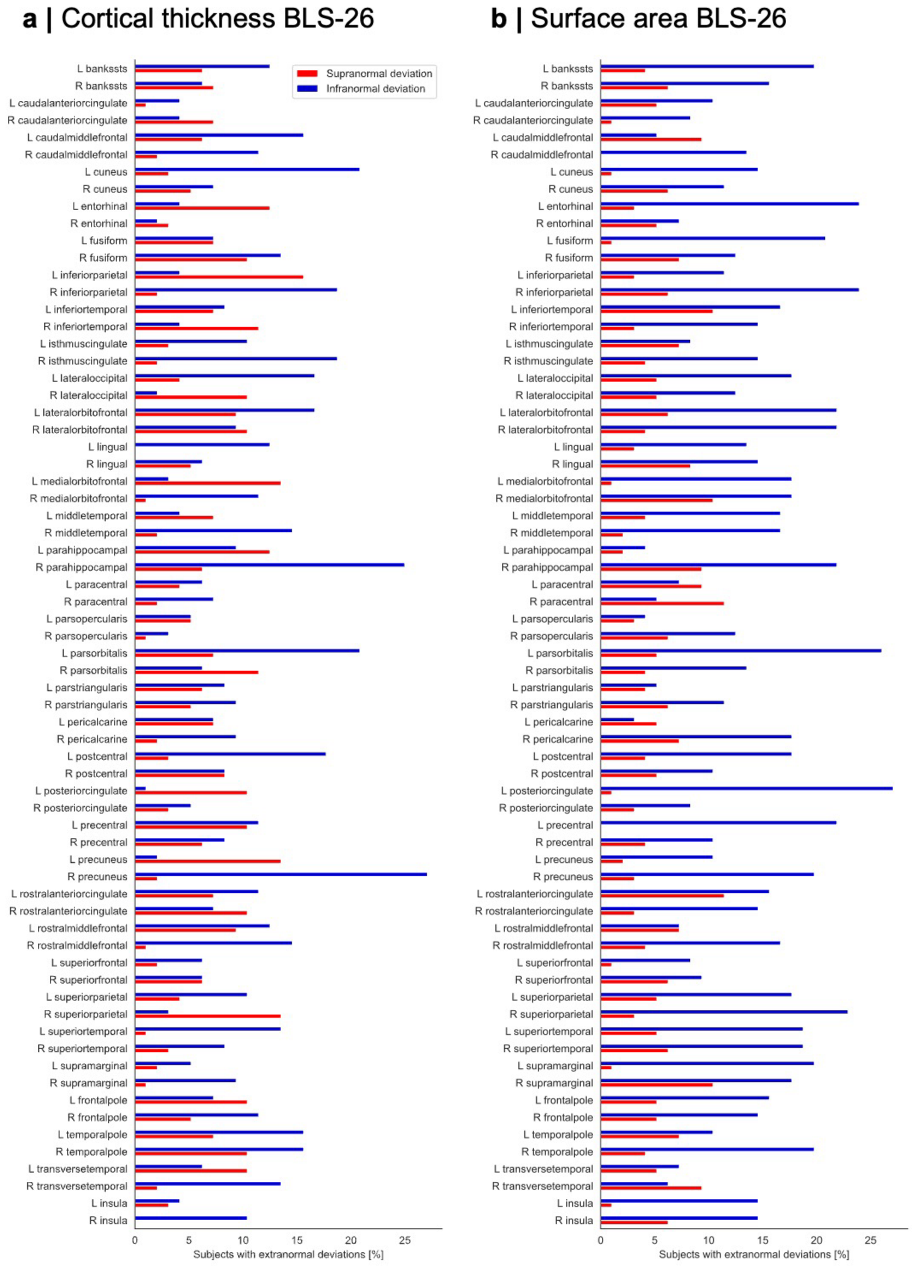

**Supplementary Figure S3: Extranormal individual deviations estimated with a different population reference normative model.** As a control analysis, a different population normative model[1] was used to estimate individual deviations for each preterm subject of the BLS-26 years cohort. Bars represent the percentage of subjects with an infranormal (i.e., < 5th percentile) or supranormal (i.e., > 95th percentile) for each region.

#### S4: Individual heterogeneity of regional surface area after preterm birth across cohorts

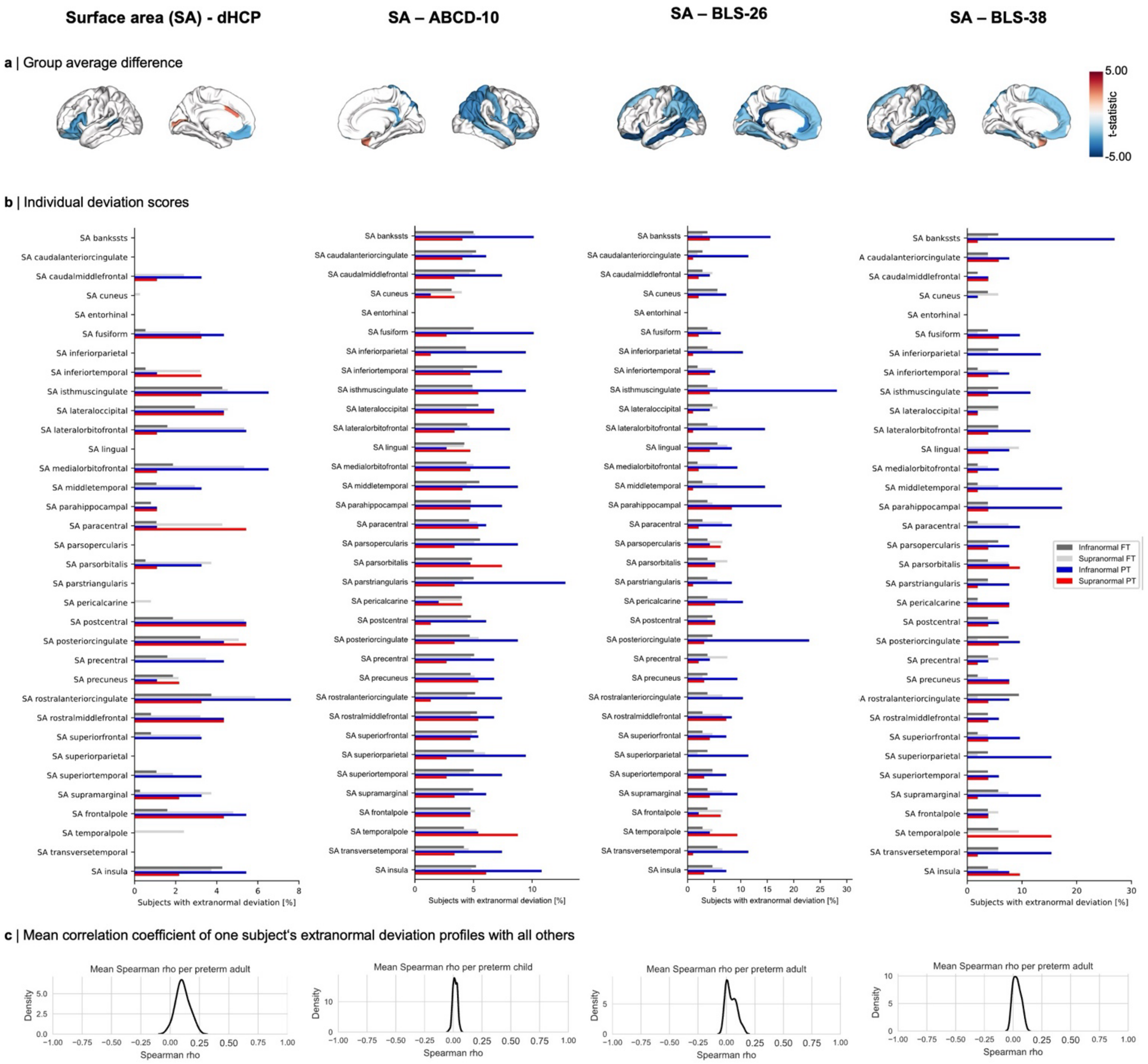

**Supplementary Figure S4: Individual heterogeneity of regional surface area after preterm birth across cohorts. a,** Surface area (SA) average dysmaturation outcome after preterm birth estimated by T-statistics corrected for age and sex ($p_{FDR} < 0.05$). **b,** The percentage of subjects sharing an extranormal deviation in any cortical region. Not more than 30 % of subjects overlap in any region, demonstrating that spatial heterogeneity between individuals is also evident for regional SA development after preterm birth. **c,** Distribution of averaged correlation coefficients of binarized extranormal deviation profiles for each subject with all others. Distributions were calculated by cross-correlating binarized extranormal deviation profiles for each individual across subjects as described in the main text. Results are shown for neonates (dHCP), children (ABCD-10), and adults (BLS-26 and BLS-38).

#### S5: Individual heterogeneity of cerebral tissue volume measures after preterm birth across cohorts

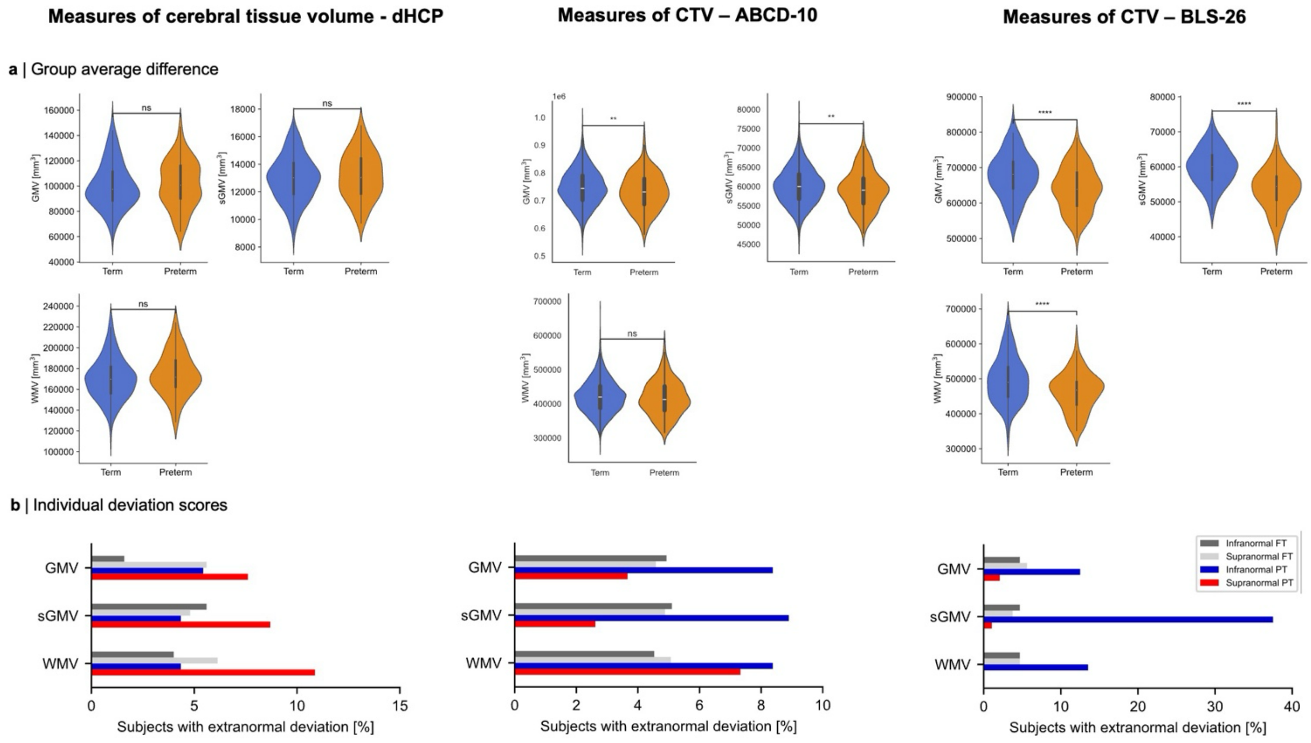

**Supplementary Figure S5: Individual heterogeneity of cerebral tissue volume measures after preterm birth across cohorts. a,** Group average difference of grey matter volume (GMV), subcortical GMV (sGMV), and white matter volume (WMV) for preterm and full-term subjects estimated by T-statistics ($p_{FDR} < 0.05$). **b,** Bar plots show the percentage of subjects sharing an extranormal deviation for the given cerebral tissue volume measure. Results are shown for neonates (dHCP), children (ABCD-10), and adults (BLS-26).

#### S6: Control analyses for the extent consistency of individual brain abnormality patterns over time

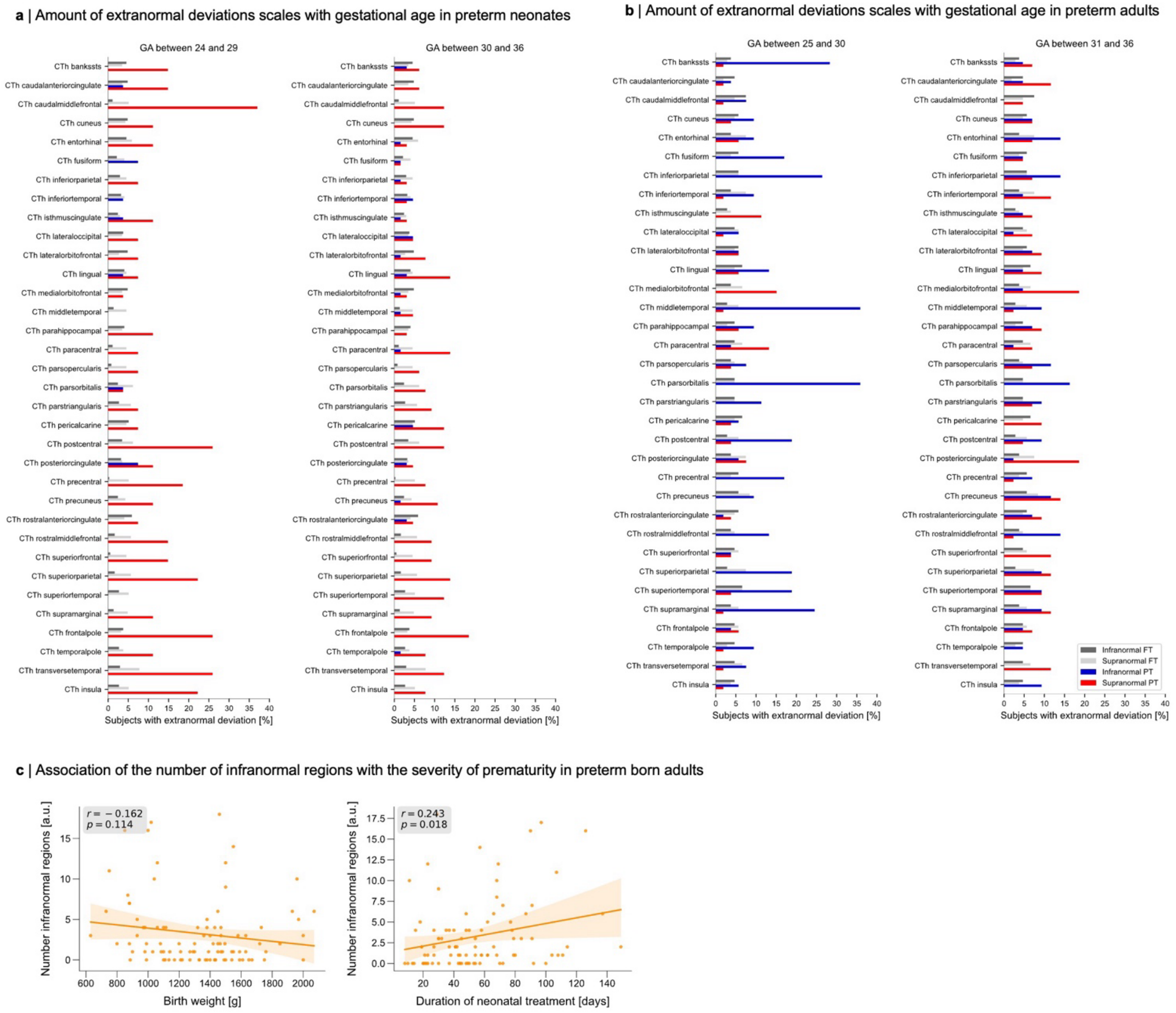

**Supplementary Figure S6: Control analyses for the extent consistency of individual brain abnormality patterns over time. a,** Preterm individuals of the dHCP cohort were divided into two groups of subjects born before (n = 27) and after (n = 65) the 30$^{th}$ week of gestation and compared the percentage of subjects that have extranormal deviations of cortical thickness (infranormal: < 5th percentile, supranormal: > 95th percentile) for any region. **b,** We divided preterm individuals of BLS-26 into two groups of earlier birth (gestational age (GA) <= 30 weeks, n = 53) and later birth (GA > 30 weeks, n = 43) and compared the percentage of subjects with infranormal (i.e., < 5$^{th}$ percentile) and supranormal (i.e., > 95$^{th}$ percentile) deviations. Whereas after earlier preterm birth, up to 36 % showed an infranormal deviation in any overlapping region, this was the case for only up to 19 % of subjects born later in the preterm period. Regions numbered from 1 to 34 correspond to the ones in panel a. **c,** We correlated the number of infranormal cortical thickness regions per subject in the BLS-26 cohort with measures of the severity of prematurity, i.e., birth weight (BW) and the Duration of Neonatal Treatment Index (DNTI). The number of infranormal regions per subject was significantly associated with DNTI (Spearman rho = 0.242, p = 0.018) but not BW (Spearman rho = -0.162, p = 0.116).

#### S7: Anatomical lesion consistency of individual deviations from 10 to 12 years

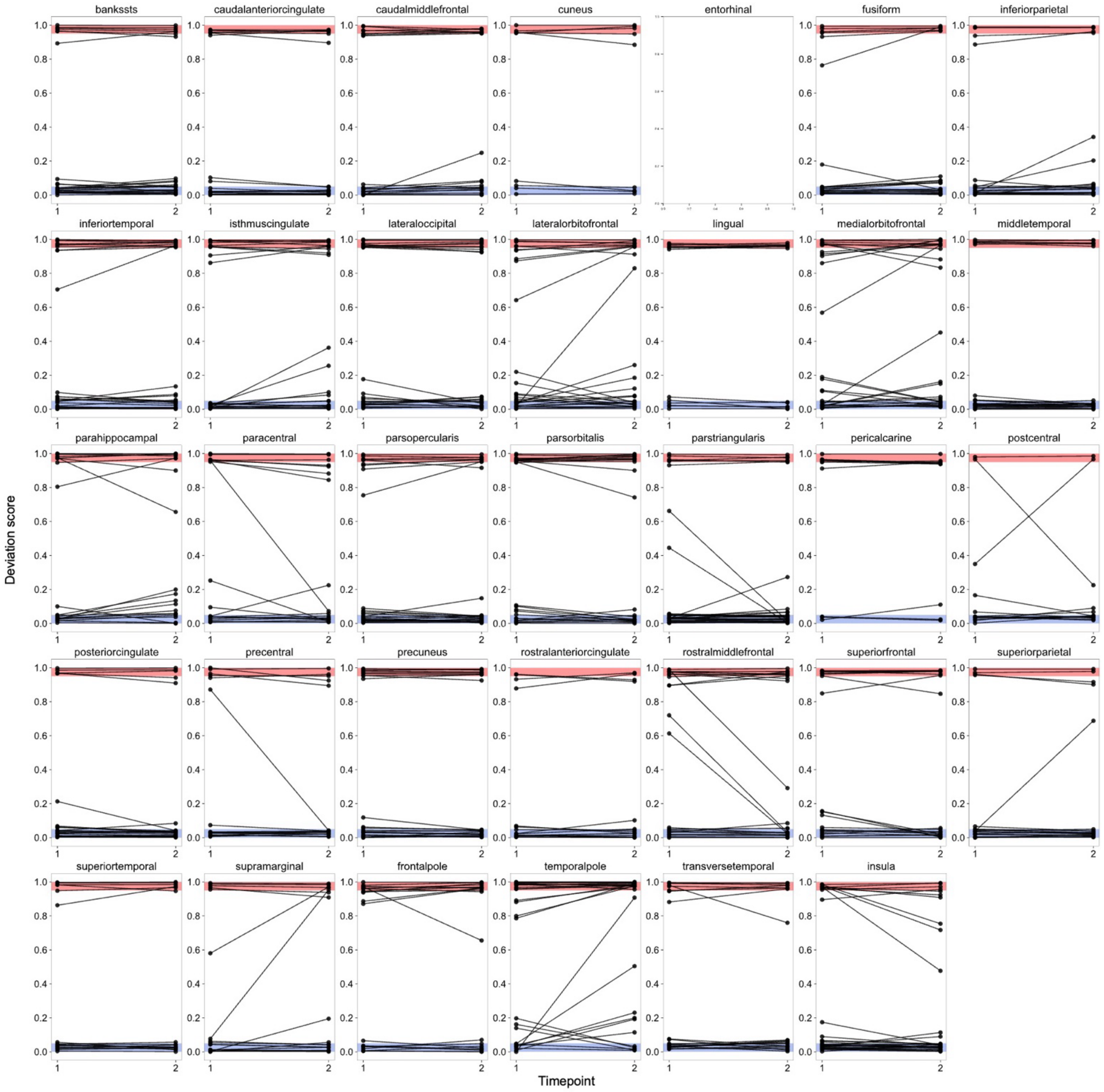

**Supplementary Figure S7: Anatomical lesion consistency of individual deviations from 10 to 12 years.** Extension to Fig. 4c. Longitudinal deviation scores of preterm children from the ABCD Study (i.e., acquired at the ages of 10 and 12 years) were calculated for regional surface area (SA) in the bilateral Desikan-Killiany parcellation. Only deviation scores of subjects that showed an extranormal deviation at either timepoint are depicted. Estimations for the entorhinal cortex were not possible due to missing information in the BrainChart reference models. Within-subject comparison between the two timepoints of data acquisition demonstrates that the anatomical location of infranormal (blue) and supranormal (red) deviations after preterm birth mostly remain consistent along childhood.

#### S8: Anatomical lesion consistency of individual deviations from 26 to 38 years

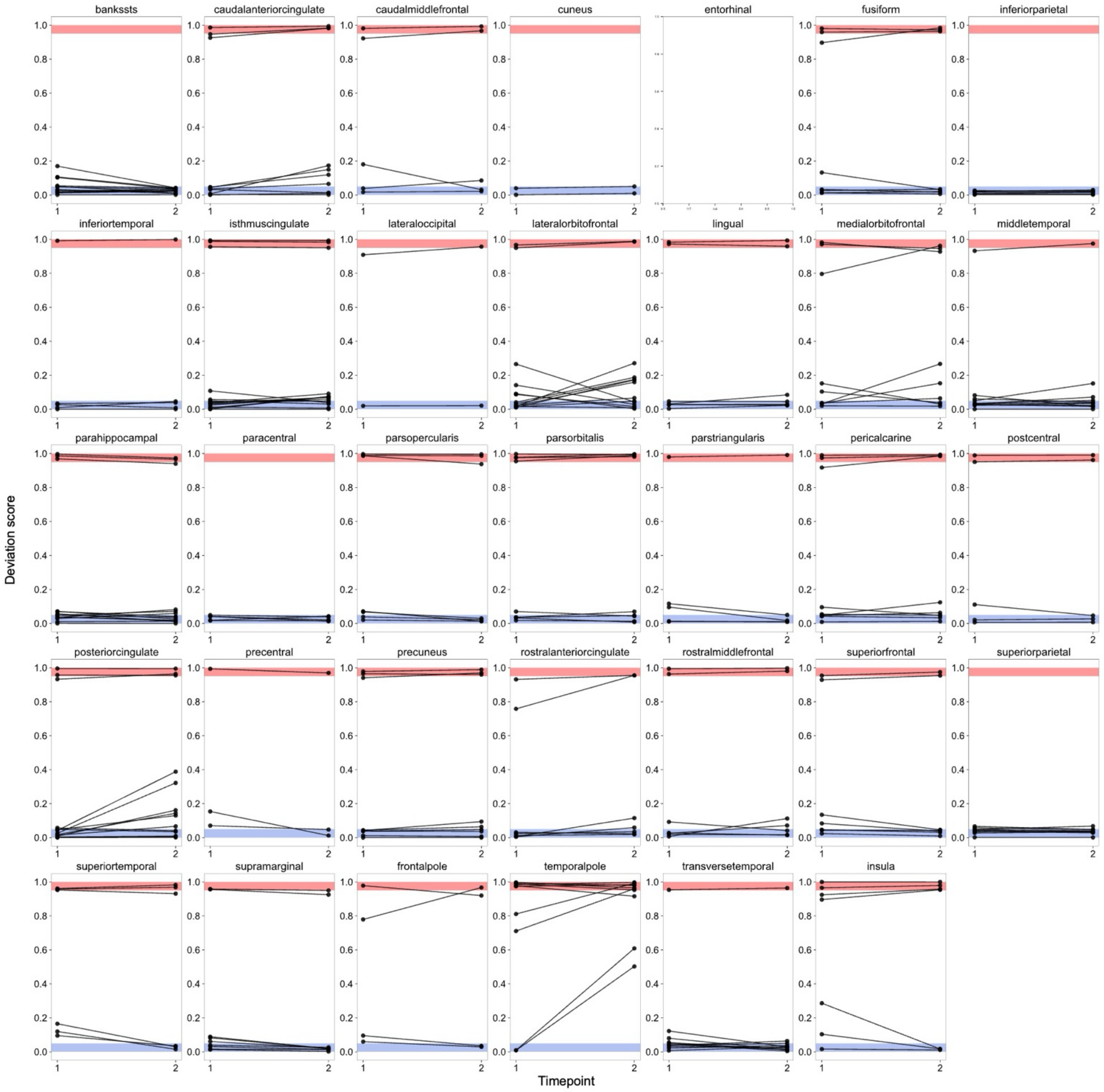

**Supplementary Figure S8: Anatomical lesion consistency of individual deviations from 26 to 38 years.** Extension to Fig. 4d. Deviation scores of preterm adults with longitudinal data (i.e., acquired at the age of 26 and partly again at the age of 38 years) were calculated for regional surface area (SA). Only deviation scores of subjects that showed an extranormal deviation at either timepoint are depicted. Estimations for the entorhinal cortex were not possible due to missing information in the BrainChart reference models. Within-subject comparison between the two timepoints of data acquisition demonstrates that the anatomical location of infranormal (blue) and supranormal (red) deviations after preterm birth mostly remain consistent along adulthood.

**S9: Significant associations between individual CTh deviations of preterm adults and mean expression profile of eight brain cell types**

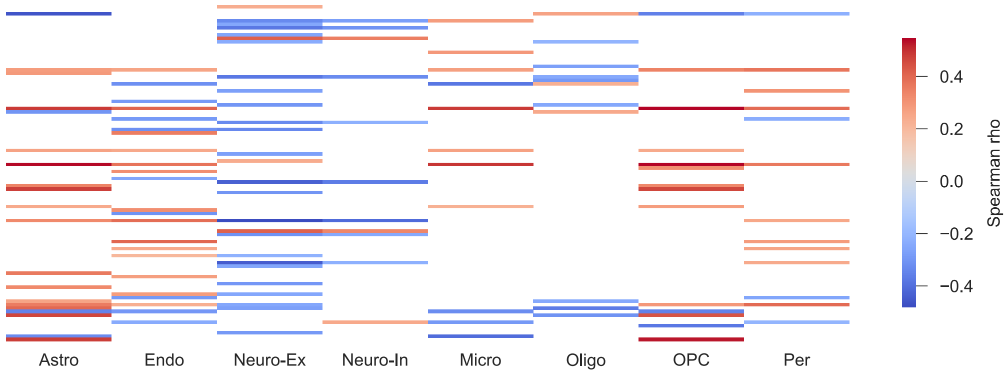

**Supplementary Figure S9: Significant associations between individual CTh deviations of preterm adults and mean expression profile of eight brain cell types**. In comparison to Fig. 5a, this Figure only shows significant associations ($p_{spin} < 0.05$). Astro astrocytes, Endo endothelial cells, Micro microglia, Neuro-Ex excitatory neurons, Neuro-In inhibitory neurons, Oligo oligodendrocytes, OPC oligodendrocyte progenitor cells, Per pericytes.

#### S10: PC loadings

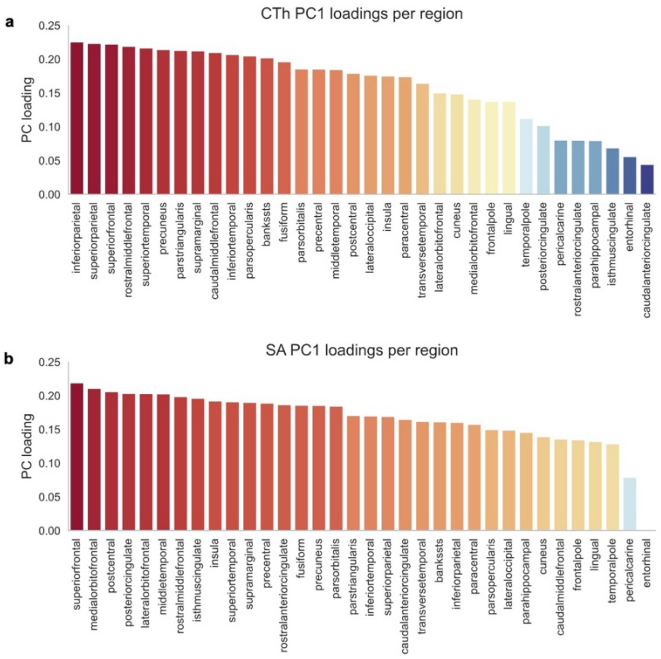

**Supplementary Figure S10: PC loadings**. Loadings of the principal component 1 from principal component analysis across 34 cortical regions for CTh (**a**) and SA (**b**). Cortical region's PC loadings are ordered from highest to lowest.

S11: Control analyses for the associations between CTh PC1 and socio-economic status

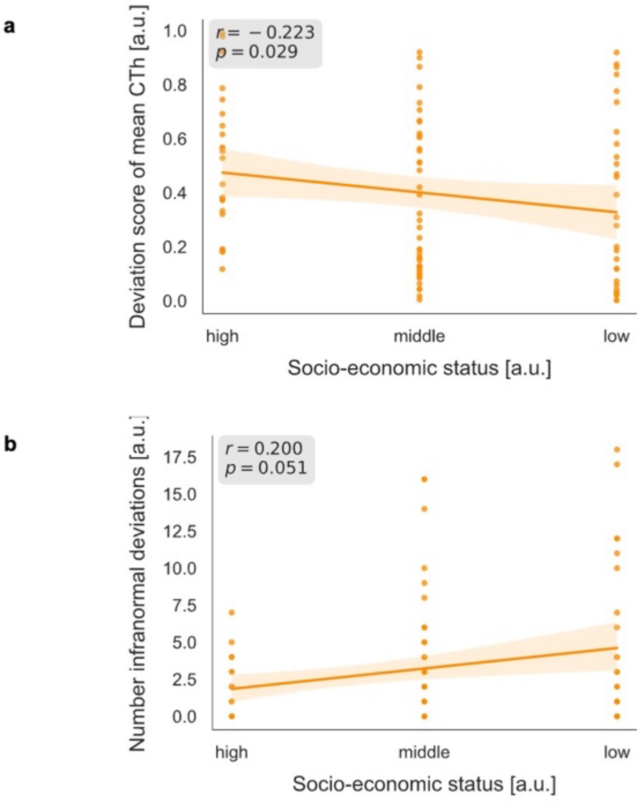

**Supplementary Figure S11: Control analyses for the associations between CTh PC1 and socio-economic status.** Instead of principal component 1 (PC1, see Fig. 3), the deviation scores of mean CT across all 34 cortical regions (**a**) as well as the number of infranormal regions per subject (**b**) were used for Spearman correlation analysis. Data of BLS-26 years preterm adults were used.

#### S12: Control analyses for the associations between CTh PC1 and IQ

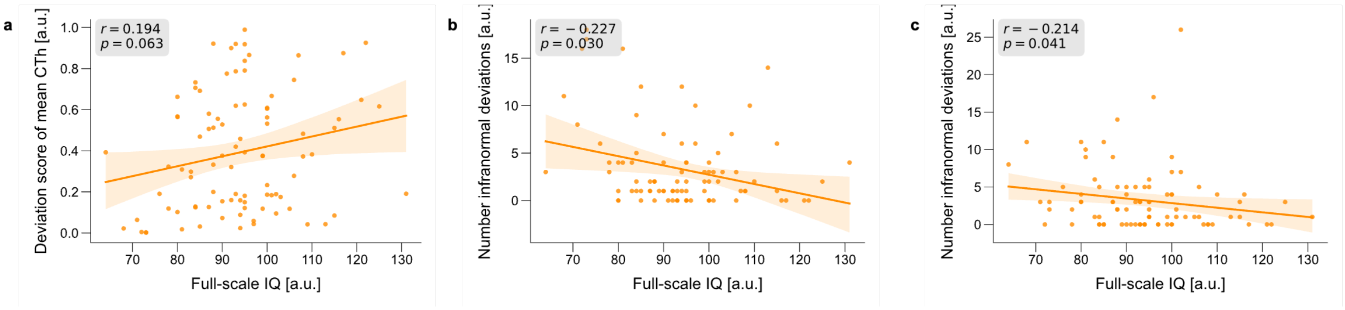

**Supplementary Figure S12: Control analyses for the associations between CTh and SA PC1 scores and IQ.** Instead of principal component 1 (PC1, see Fig. 4), the deviation scores of mean CTh across all 34 cortical regions (**a**) as well as the number of infranormal regions per subject (**b**) were used for Spearman correlation analysis. Furthermore, the number of infranormal regions of regional SA were correlated with IQ measures (**c**). Data of BLS-26 years preterm adults were used.

#### Supplementary tables

##### Table S1: Demographics

***Table 1: Demographical, clinical, and cognitive data**. Data are listed for the final data sample after exclusion due to quality control.*

The table displays mean ± SD, except where noted.

*Statistical comparisons*: sex and SES with $\chi^2$ statistics; age, GA, BW, full-scale IQ with two-sample t-tests. Bold letters indicate statistical significance defined as $p < 0.05$.

*Abbreviations*: BW, birth weight; FT, full-term; GA, gestational age; IQ, intelligence quotient; n.a., not applicable; SD, standard deviation; SES, socio-economic status; PT preterm.
